# HESTIA: Scalable Multimodal Integration of Histology and High-Resolution Spatial Transcriptomics for Robust Spatial Domain Identification

**DOI:** 10.64898/2026.05.14.723098

**Authors:** Zheng Zhong, Xiaoyu Zhu, Jing Guo, Sha Liao, Ao Chen

## Abstract

Spatial omics has revolutionized molecular biology by providing invaluable insights into how native tissue microenvironments regulate cellular functions and disease mechanisms. Accurately capturing this structural complexity and decoding the underlying biological processes requires effectively integrating data from multiple modalities. However, transitioning to subcellular resolutions introduces massive data scales and severe transcriptomic sparsity, which challenge current analytical frameworks. To address this, we present HESTIA (Histology-Enhanced Scalable cross-Resolution inTegration for spatial trAnscriptomics), a highly efficient multimodal algorithm designed for identifying spatial domains in large-scale, high-resolution spatial omics data. By circumventing memory-intensive computations, HESTIA efficiently processes massive datasets on which existing algorithms fail due to memory constraints. HESTIA outperforms current multimodal methods in clustering accuracy and spatial continuity, accurately delineating fine structural boundaries. Furthermore, applying HESTIA to large-scale pathological samples successfully dissects clinically relevant intratumoral heterogeneity and maps distinct immune microenvironments in lung and colorectal cancers.

## Introduction

Spatial transcriptomics has revolutionized molecular biology by anchoring gene expression profiles to their exact physical coordinates within tissue structures^1^. Unlike single-cell RNA sequencing (scRNA-seq), which requires tissue dissociation and consequently loses spatial context, spatial transcriptomics preserves the intricate spatial relationships among cells. This *in situ* perspective is indispensable for decoding cell-cell communications and understanding how distinct cellular states are modulated by their native microenvironments^2^. Furthermore, most spatial transcriptomics platforms concurrently acquire high-resolution morphological data, such as hematoxylin and eosin (H&E) stained histology images. These visual profiles have long served as the gold standard for clinical decision-making, particularly in oncology^3^. Leveraging these paired histological images as an orthogonal data modality allows researchers to cross-validate observations, counteract technical noise, and compensate for the dropout events inherent to pure RNA-seq data^4^.

A primary objective in spatial transcriptomics analysis is the robust characterization of spatial domains. These spatial domains emerge as higher-order structural units composed of coordinated cell populations exhibiting distinct transcriptional signatures^5^. Two complementary paradigms exist for this task. While the cell-centric paradigm defines domains by first segmenting and annotating cells and then assembling them into niches based on cellular composition, an alternative paradigm partitions tissue directly from the expression and morphological features of spatial units^6^. This work focuses on the latter, motivated by its independence from cell type annotations, its robustness to cell segmentation errors, and its ability to capture tissue structures through both molecular and morphological cues. Following this direct paradigm, recent methods integrate morphological imagery with spatial gene expression, providing complementary information to identify heterogeneities that might be missed by single-modality analyses. For instance, SpaGCN^7^ employs graph convolutional networks to harmonize gene expression, spatial coordinates, and imaging features. MUSE^8^ utilizes a multimodal autoencoder and triplet loss to integrate cellular morphologies with transcriptional states. Similarly, ConGR^9^ uses contrastive learning and graph autoencoder to align distinct data modalities, and StereoMM^10^ introduces a graph fusion model bolstered by cross-modal attention mechanisms.

However, a critical limitation of these existing methods stems from their development and evaluation on early-generation spatial transcriptomic datasets characterized by lower resolution or restricted throughput. Modern platforms generate data at unprecedented scales: Visium HD^11^ offers resolution as refined as 2 μm, and Stereo-seq^12^ can achieve up to 0.5 μm resolution across expansive slice areas of up to 13 cm × 13 cm. Advances in resolution introduce two unique challenges. First, the steep increase in data volume causes existing algorithms, which rely on computationally intensive graph neural networks or attention modules, to suffer from severe time and/or memory bottlenecks^13,14^. Second, scaling down to subcellular resolutions exacerbates transcriptomic sparsity, resulting in higher rates of gene dropout and incomplete molecular localization^15^.

To bridge this gap, we propose HESTIA (Histology-Enhanced Scalable cross-Resolution inTegration for spatial trAnscriptomics), a highly scalable, multimodal algorithm designed for the high-resolution spatial transcriptomics era. HESTIA integrates multiscale tissue morphology and cross-resolution transcriptomic features to drive accurate spatial domain identification. Rather than relying on computationally intensive architectures, HESTIA employs a multimodal learning framework that integrates histology and transcriptomic data. A dual-autoencoder with cross-resolution alignment helps mitigate transcriptomic sparsity, while the fusion of morphological and molecular features ensures robust and scalable spatial domain identification.

We demonstrate that HESTIA successfully processes the full-slice Stereo-seq mouse brain dataset, a scale at which competing algorithms critically fail due to memory constraints. Quantitative assessments of subregions in this mouse brain dataset reveal that HESTIA consistently achieves superior clustering accuracy and spatial coherence compared to existing multimodal algorithms. Furthermore, we demonstrate HESTIA’s practical applicability by successfully delineating tumor subregions in a Stereo-seq human lung adenosquamous carcinoma sample and a Visium HD human colorectal cancer dataset. By overcoming the challenges posed by extreme data scale and feature sparsity, HESTIA offers an invaluable analytical framework for unlocking the full biological potential of high-resolution spatial multi-omics.

## Results

### Overview of HESTIA

HESTIA is a multimodal analytical framework that integrates high-resolution spatial transcriptomic data with paired H&E histology images via an efficient, multiscale learning architecture (Figure 1A). To capture morphological patterns, HESTIA employs a pre-trained Hierarchical Vision Transformer (HViT)^16^ to extract global and local structural representations from high-resolution histological images. These representations are then processed by an autoencoder to generate image features that are dimensionally aligned with the spatial transcriptomic features. Recognizing that neighboring locations on tissue sections often share cellular microenvironments and exhibit similar gene expression profiles, HESTIA applies a spatial binning strategy to generate a parallel, low-resolution transcriptomic representation. This spatial aggregation also effectively mitigates the signal sparsity inherent in high-resolution spatial transcriptomics platforms. After preprocessing and principal component analysis (PCA), both the high-resolution and low-resolution expression matrices are fed into separate autoencoders. During training, the framework optimizes a joint loss function that combines standard reconstruction loss with cross-resolution contrastive loss. This dual-autoencoder design enforces representation consistency between high- and low-resolution transcriptomic states, helping to stabilize sparse molecular signals. The resulting transcriptomic features are then fused with histology-derived features to produce a robust multimodal representation, which provides a solid foundation for spatial domain identification. Finally, to maximize computational efficiency during practical use, HESTIA incorporates a modular caching mechanism that stores intermediate feature outputs. When algorithmic parameters are adjusted, the framework directly reuses unaffected upstream results, which significantly reduces overall runtime and computational overhead.

**Figure 1.**
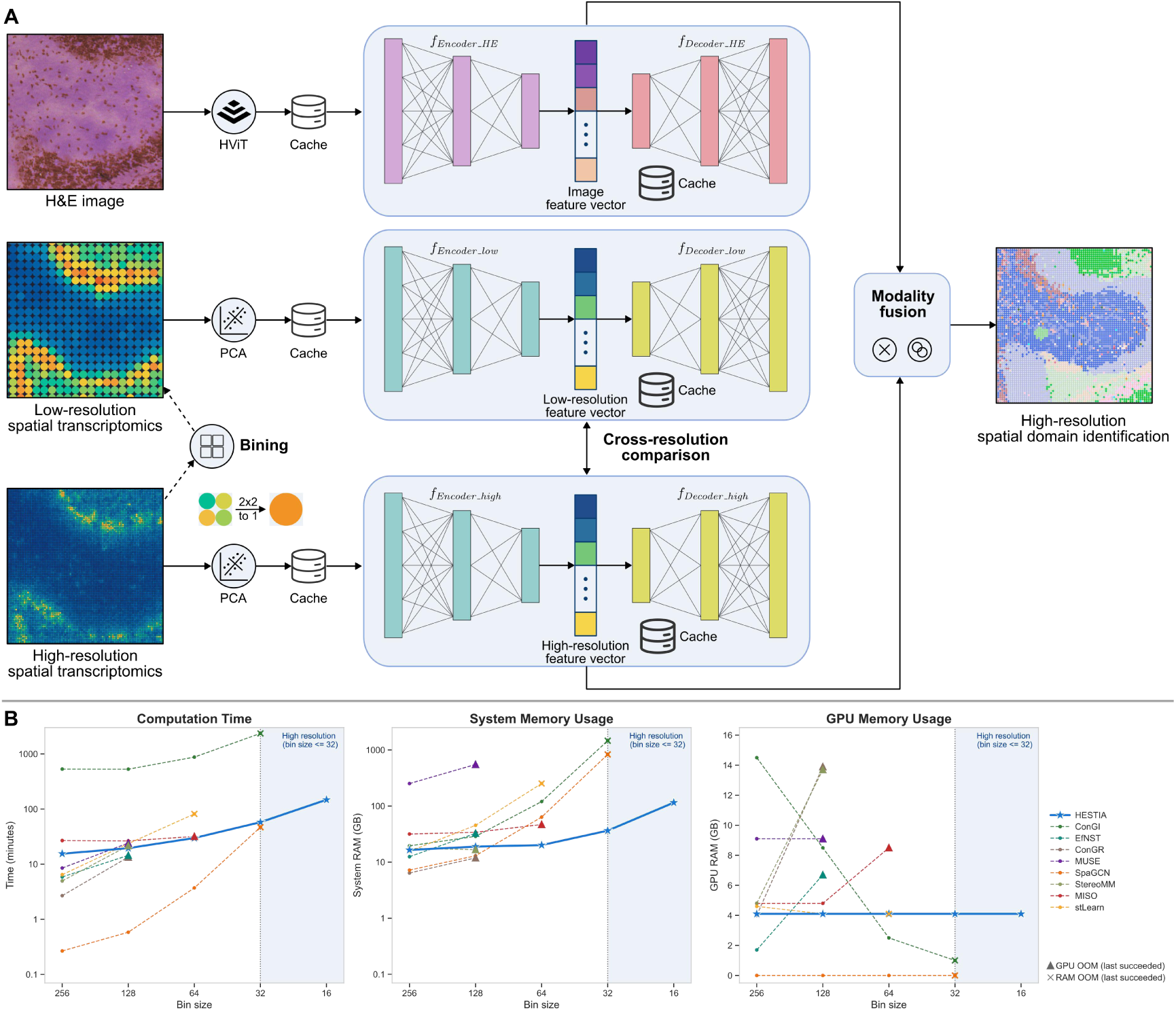
Overview of HESTIA and evaluation of computational scalability. (A) Schematic representation of the HESTIA architecture. HESTIA uses co-registered high-resolution spatial transcriptomics data and H&E histology images as inputs. Top branch: Histological representations are extracted using a pre-trained HViT model and compressed into an image feature vector via an image autoencoder. Middle and bottom branches: To mitigate extreme signal sparsity, the high-resolution spatial transcriptomic data are spatially aggregated (e.g., 2×2 binning) to generate a parallel, low-resolution transcriptomic profile. After PCA, both the high- and low-resolution spatial transcriptomic data are processed through a dual-autoencoder system. A cross-resolution comparison is enforced between their latent spaces to stabilize the sparse molecular signals. Modality fusion: The resulting high-resolution transcriptomic feature vector and the image feature vector are integrated using outer product and concatenation operations. The fused multimodal representation is then utilized for high-resolution spatial domain identification. A modular caching system stores intermediate outputs at multiple steps, drastically reducing redundant computations during parameter tuning. (B) Benchmarking the computational efficiency and scalability of HESTIA and other multimodal algorithms. The shaded blue area represents the high-resolution threshold (bin size ≤ 32). Triangles and crosses indicate the last successful bin size at which an algorithm was able to complete before failing due to GPU or system RAM out-of-memory (OOM), respectively.

### HESTIA exhibits high computational efficiency and scalability

We evaluated the computational efficiency of HESTIA alongside eight other multimodal methods: ConGI^17^, ConGR^9^, EfNST^18^, MISO^19^, MUSE^8^, SpaGCN^7^, StereoMM^10^, and stLearn^20^. For this benchmark, we used a high-resolution Stereo-seq mouse brain dataset. A key advantage of using Stereo-seq data is the ability to easily apply different bin sizes to obtain varying numbers of data points (bins) from a single dataset while maintaining consistency in all other biological and technical variables. Specifically, we processed the data using bin sizes ranging from bin256 to bin16. The highest resolution bin16 contains over 818,000 bins, reflecting the large scale of modern high-resolution spatial transcriptomics platforms (Supplementary Table 1).

As the bin size decreased, the increased data volume exposed significant computational bottlenecks in all tested methods. ConGR, EfNST, MUSE, and StereoMM could not complete the bin64 analysis with 16 GB of GPU RAM, while MISO failed the bin32 analysis with the same 16 GB GPU limit. stLearn could not complete the bin32 analysis, and both ConGI and SpaGCN failed the bin16 analysis, even with 2000 GB of system RAM allocated (Figure 1B). In contrast, HESTIA was the only method that successfully analyzed the massive bin16 data with reasonable memory consumption. HESTIA completed the bin16 analysis in under 120 minutes using less than 120 GB of system memory and approximately 4 GB of GPU memory, demonstrating its highly scalable nature (Figure 1B). We note that this efficiency gain partly reflects a deliberate architectural trade-off: HESTIA bypasses the explicit spatial graph construction that underlies most competing methods and constitutes their primary memory bottleneck. Additionally, enabling caching would allow the tissue image to be processed only once for different bin sizes, saving even more computational resources (Supplementary Figure 1).

### HESTIA accurately identifies spatial domains in high-resolution Stereo-seq mouse brain dataset

We evaluated the spatial domain identification performance of HESTIA using the same Stereo-seq mouse brain dataset because mouse brain has clear hierarchical structures. To ensure that all algorithms could analyze the data without encountering memory issues, we subset the data into seven distinct subregions. Each subregion contains approximately 10,000 bins at the bin20 resolution and covers the cortex, hippocampus, striatum and thalamus/hypothalamus (Supplementary Figure 2A). We manually annotated these subregions based on reference annotations from the Allen Mouse Brain Atlas^21^ (Figure 2A and Supplementary Figure 3). To validate the accuracy of manual annotations, we performed cell2location^22^ annotation using scRNA-seq data from the Allen Brain Cell Atlas^23^. The resulting cell-type maps show high concordance with our structural annotations (Supplementary Figure 2B, C). To quantify spatial domain identification performance, we assessed clustering accuracy using the adjusted Rand index (ARI) and normalized mutual information (NMI), and spatial continuity using average silhouette width (ASW)^24^, CHAOS^25^, and percentage of abnormal spots (PAS)^25^. Higher values of ARI, NMI, and ASW indicate better spatial domain identification performance, whereas lower values of CHAOS and PAS reflect better spatial coherence.

Visual inspection of an example mouse brain subregion demonstrated that HESTIA accurately identified the complex structure of hippocampus with clear boundaries. Notably, the domain assignments generated by HESTIA were highly contiguous and contained almost no scattered spots (Figure 2B). Evaluation metrics confirmed these visual observations, showing HESTIA consistently achieved the best clustering accuracy and continuity scores across all seven subregions (Figure 2D). stLearn and SpaGCN identified spatial domains similar to HESTIA, but their predictions had lower overall accuracy and contained more dispersed spots, indicating reduced spatial continuity (Figure 2C,D). The remaining multimodal algorithms—including MISO, ConGI, ConGR, EfNST, StereoMM, and MUSE—struggled to define the boundaries of the hippocampus and the distinct layers of the cortex. These methods produced noisy cluster assignments with high degrees of mixing between anatomically distinct zones, corresponding to their lower ARI, NMI, and ASW scores, as well as their elevated CHAOS and PAS values, across the seven subregions in this high-resolution dataset (Figure 2C, D). Our primary benchmark focused on multimodal methods, as HESTIA is designed to integrate histology and transcriptomics. For comprehensiveness, we also compared HESTIA with three scalable single-modality methods that do not utilize image features: BINARY^26^, MENDER^27^, and BANKSY^28^. Although these methods are also computationally efficient, their spatial domain assignments were less accurate than HESTIA (Supplementary Figures 3, 4). These results further support the advantage of integrating histological information for spatial domain identification.

**Figure 2.**
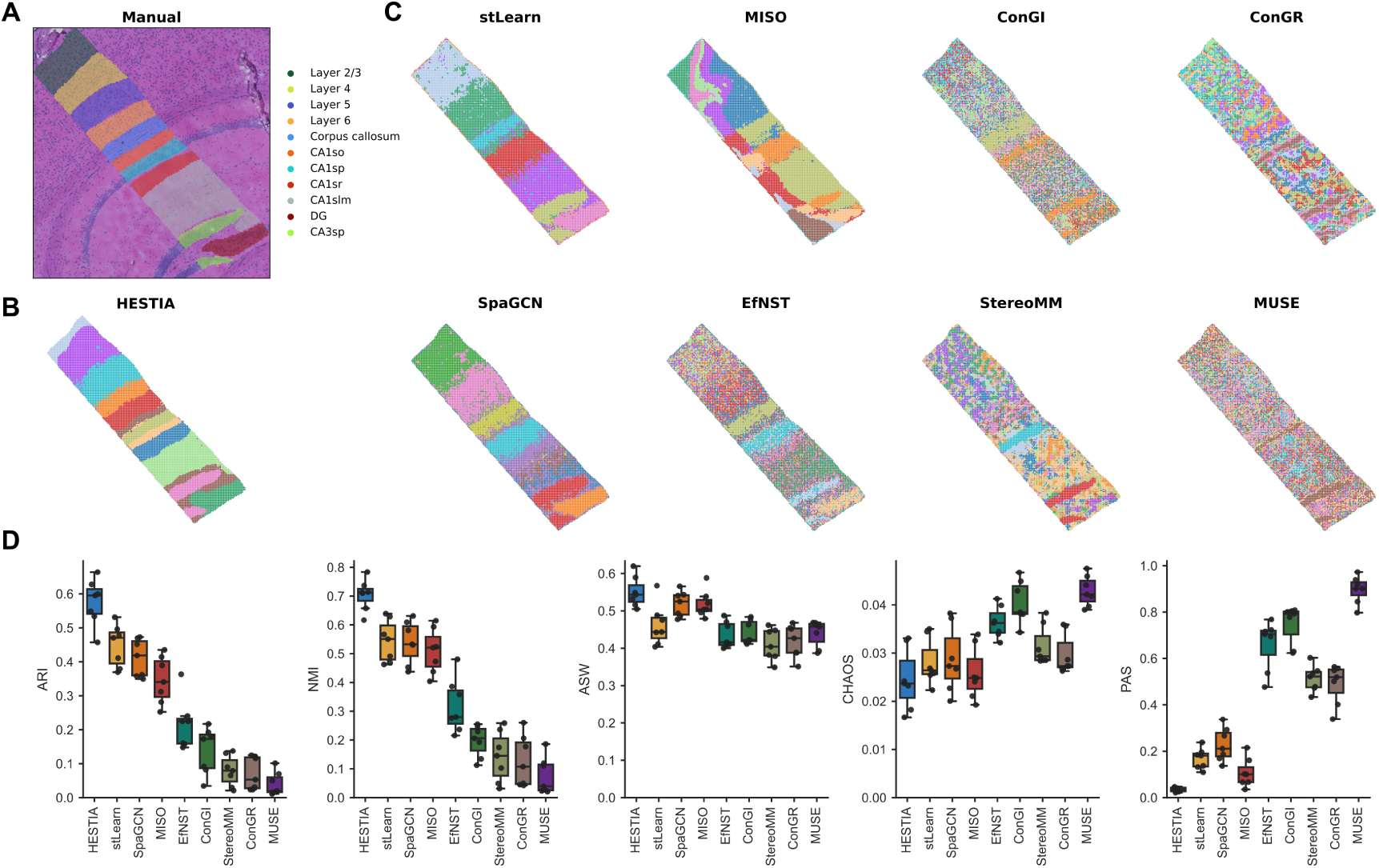
Performance benchmark of spatial domain identification in high-resolution Stereo-seq mouse brain data (bin20 level). (A) H&E-stained histology image of a representative subregion spanning the mouse brain cortex and hippocampus, overlaid with manual annotations based on the Allen Mouse Brain Atlas. (B) Spatial domain identification results generated by HESTIA for the representative tissue subregion shown in (A). (C) Spatial domain identification results generated by other multimodal algorithms applied to the same tissue subregion. (D) Quantitative evaluation of spatial domain identification performance across seven mouse brain subregions. Boxplots display the clustering accuracy metrics (ARI and NMI) and the spatial continuity metrics (ASW, CHAOS, and PAS) for HESTIA and other algorithms. Higher ARI, NMI, and ASW scores denote superior performance, whereas lower CHAOS and PAS values indicate greater spatial coherence. The center lines of the boxplots represent the median, the box boundaries represent the interquartile range (IQR), and the whiskers extend to 1.5x IQR.

### HESTIA enables highly precise spatial domain identification at single-cell resolution

After successfully applying HESTIA to grid-based spatial data, we evaluated its capacity to resolve tissue structure at true single-cell resolution. In the Stereo-seq platform, spatial transcriptomic data are obtained by detecting unique molecular identifiers captured by DNA nanoballs (DNBs) patterned on the sequencing chip with a highly granular 0.5 μm resolution (bin1)^12^. By leveraging paired histology images, cellular segmentation can aggregate these distinct subcellular signals into individual single-cell spatial profiles^29^. We used the single-cell version (cellbin) of the aforementioned Stereo-seq mouse brain dataset, which was segmented via the Stereo-seq Analysis Workflow (SAW)^30^ to yield approximately 140,000 individual cells. We first applied HESTIA to the full-slice mouse brain cellbin data (Figure 3A). Compared to the coarser bin-level analyses (Supplementary Figure 5), the spatial domains identified in the cellbin data exhibited a markedly finer separation of distinct anatomical structures (Figure 3B). Notably, the spatial domains defined by HESTIA showed a visual concordance with the established anatomical annotations from the Allen Mouse Brain Atlas (Figure 3C).

Given that processing a dataset of 140,000 cells severely exceeds the memory limits of most alternative tools, we restricted our comparative evaluation with other algorithms to the seven subregions which we used earlier. Visual examination of HESTIA’s output confirmed its ability to delineate complex and fine-grained structures accurately. It outlined the distinct cortical layers, the corpus callosum, the dentate gyrus (DG), and the stratum pyramidale (sp) of both the cornu ammonis 1 (CA1) and CA3, as well as the CA1 stratum lacunosum-moleculare (slm) (Figure 3D, E, G, H).

When applied to the same subregions, several existing multimodal algorithms encountered difficulties in fully resolving these subtle structural boundaries. For instance, MISO struggled to isolate specific structural tracts, such as the corpus callosum and CA3sp. While stLearn and SpaGCN outlined the broader major anatomical regions, they displayed minor boundary overlaps in their spatial assignments, with some scattered spots blending into adjacent domains. EfNST and ConGI tended to group distinct cortical layers into broader, overlapping domains. Similarly, StereoMM, ConGR, and MUSE produced domain assignments across the cortex and hippocampus that were intermixed and lacked defined biological boundaries (Supplementary Figure 6).

To validate HESTIA’s superior structural demarcations, we mapped the spatial distribution of canonical marker genes against our computational assignments. Consistent with our domain predictions, the expression of the CA1sp marker gene *Wfs1* was enriched within the HESTIA-annotated CA1sp territory. Similarly, the DG-specific marker gene *C1ql2* exhibited concentrated expression within the algorithmically defined DG domain. This spatial and transcriptional alignment was equally evident in *Chgb* mapping to the CA3sp region, as well as in *Mbp* localizing to the corpus callosum (Figure 3F, I).

HESTIA’s superior visual performance was confirmed through quantitative evaluation of spatial clustering metrics. In accordance with our observations on the bin20 data, HESTIA outperformed all other evaluated algorithms on the cellbin data. It achieved the highest ARI, NMI, and ASW scores while recording the lowest CHAOS and PAS scores (Figure 3J). Similarly, the single-modality methods BINARY, MENDER, and BANKSY also underperformed HESTIA at the cellbin level (Supplementary Figures 6, 7). These metrics underscored HESTIA’s exceptional clustering accuracy and spatial continuity at a single-cell resolution.

**Figure 3.**
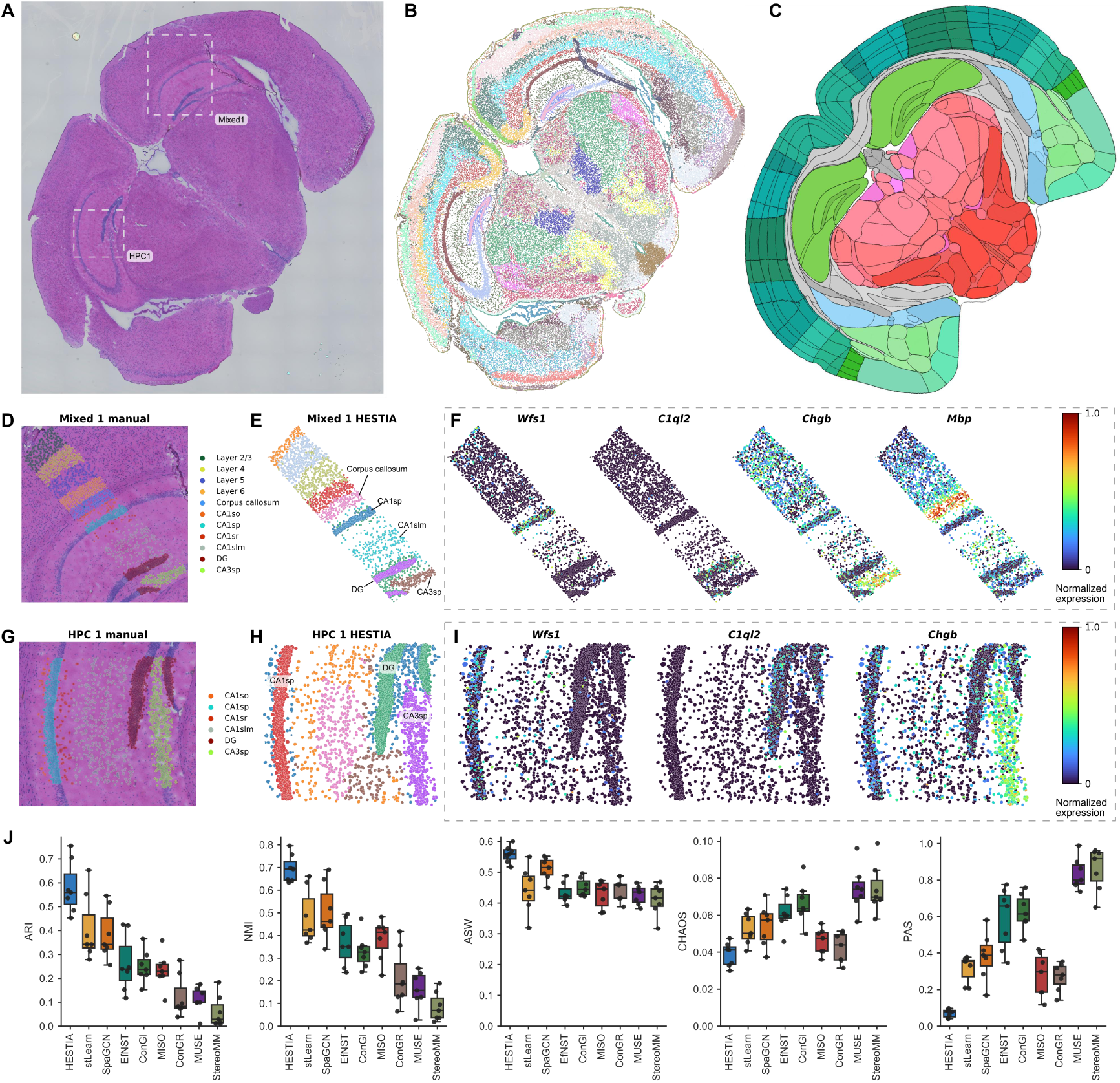
HESTIA enables highly precise spatial domain identification at single-cell (cellbin) resolution. (A) H&E-stained histology image of a full Stereo-seq mouse brain coronal section. Dashed boxes indicate the locations of two selected subregions, designated as “Mixed 1” and “HPC 1”. (B) Spatial domains identified by HESTIA across the entire coronal section at cellbin level. (C) Corresponding anatomical reference map adapted from the Allen Mouse Brain Atlas. (D) Manual spatial domain annotations of the “Mixed 1” subregion overlaid on the H&E image. (E) Spatial domains identified by HESTIA in the “Mixed 1” subregion. (F) Spatial gene expression plots of marker genes within the “Mixed 1” subregion: *Wfs1* (CA1sp), *C1ql2* (DG), *Chgb* (CA3sp), and *Mbp* (corpus callosum). (G) Manual spatial domain annotations of the “HPC 1” subregion overlaid on the H&E image. (H) Spatial domains identified by HESTIA in the “HPC 1” subregion. (I) Spatial gene expression plots of marker genes within the “HPC 1” subregion: *Wfs1* (CA1sp), *C1ql2* (DG) and *Chgb* (CA3sp). (J) Quantitative evaluation of spatial domain identification at cellbin level across seven distinct subregions. Boxplots display clustering accuracy (ARI, NMI) and spatial continuity (ASW, CHAOS, PAS) for HESTIA and the compared algorithms. Higher ARI, NMI, and ASW scores denote superior performance, whereas lower CHAOS and PAS values indicate greater spatial coherence. The center lines of the boxplots represent the median, the box boundaries represent the IQR, and the whiskers extend to 1.5× IQR.

### HESTIA characterizes distinct immune microenvironments in a massive lung adenosquamous carcinoma dataset

To examine the performance of HESTIA in more complex, clinically relevant pathological samples, we analyzed a human lung adenosquamous carcinoma dataset generated using the Stereo-seq platform^31^. The tissue section was previously annotated by pathologists into four main regions: adenocarcinoma (AC), squamous cell carcinoma (SCC), a mixed AC and SCC region, and other regions (Figure 4A). At a resolution of bin20, this dataset contains more than 2 million bins. No other tested multimodal method could analyze the full tissue section at this massive scale due to severe memory constraints. Remarkably, however, HESTIA successfully processed the entire full-slice bin20 dataset using a single GPU with 24 GB of RAM, further underscoring its scalability for modern spatial transcriptomic data.

As noted in the original study, the mixed region in this sample contains intermingled AC and SCC cells and lacks a discrete transcriptional identity. We therefore did not increase the number of clusters to force its separation. This mixed region could hardly be identified as a distinct cluster by the original study, StereoMM, or the alternative clustering approaches we tested (K-means, Leiden, and Louvain on both expression PCA and cell2location annotation scores, with varying parameter settings; Supplementary Figure 8). Under these conditions, HESTIA identified the AC, SCC, and other regions with an ARI of 0.33, an NMI of 0.35, and an ASW of 0.49 (Figure 4B, E). These results are notably better than those obtained using a single modality (Figure 4C–E), demonstrating the benefit of integrating multimodal data for spatial domain identification.

When further increasing the clustering number, HESTIA identified subdomains within the SCC domain, designated as domain 2 and domain 0 (Figure 4F). Analysis of differentially expressed genes (DEGs) showed that domain 2 was characterized by a significantly high expression of immunoregulatory genes, including *CCL19*, *IGLC2*, *IGLC3*, and *MZB1* (Figure 4G). Pathway enrichment analysis revealed that these genes are strongly associated with the immune response (Figure 4H). Gene Set Enrichment Analysis (GSEA) confirmed that domain 2 is characterized by the upregulation of B-cell immunity (Supplementary Figure 9). Matrix plots and spatial expression maps of these signature genes exhibited distinct localization patterns within domain 2 (Figure 4I, J), suggesting that HESTIA’s separation of domain 2 and domain 0 is biologically meaningful.

**Figure 4.**
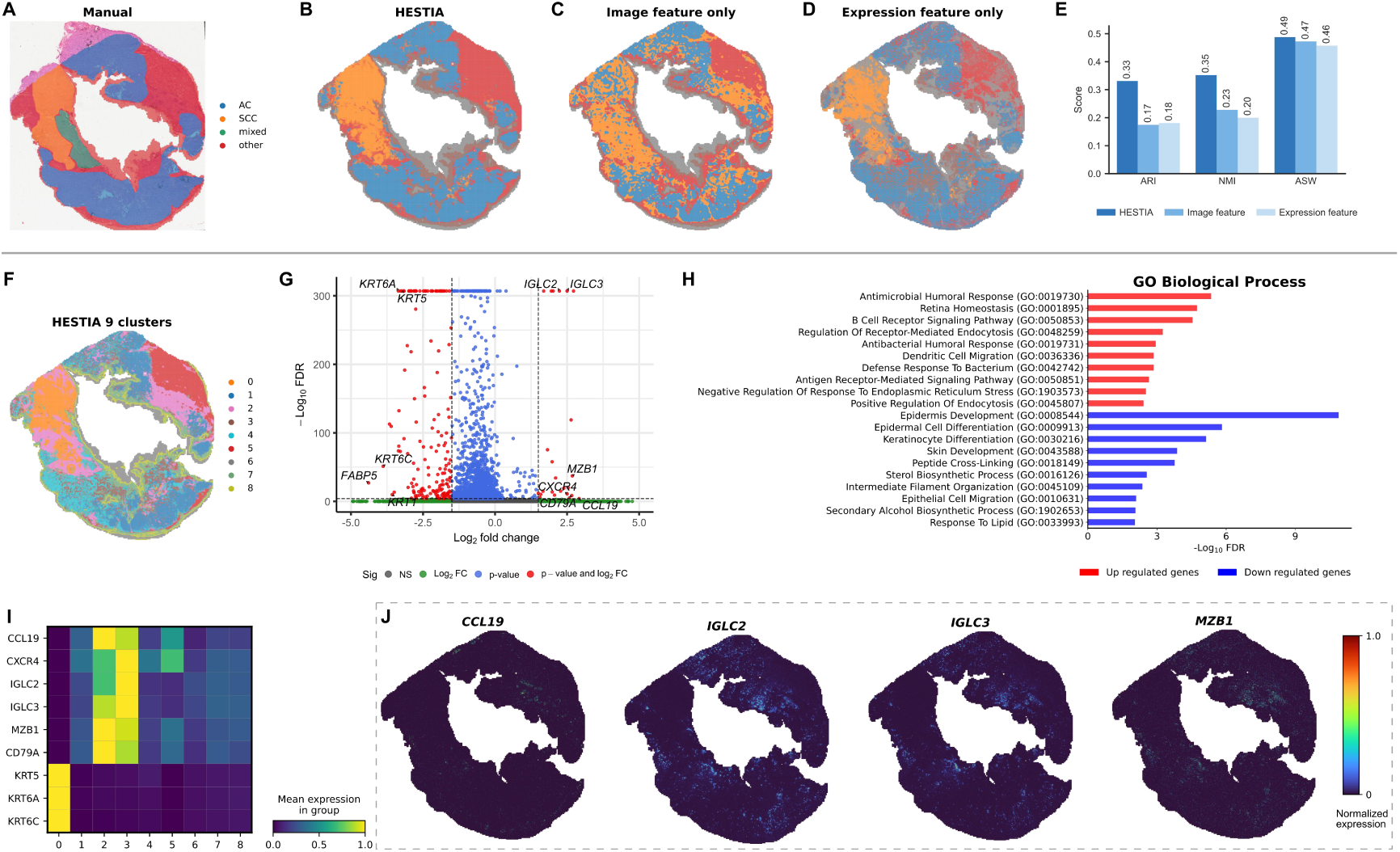
HESTIA characterizes distinct immune microenvironments in a massive lung adenosquamous carcinoma dataset. (A) Manual annotations of a full-slice human lung adenosquamous carcinoma sample. Major functional zones include adenocarcinoma (AC), squamous cell carcinoma (SCC), a mixed region, and other tissue types. (B) Spatial domain assignments generated by the full multimodal HESTIA framework. (C–D) Spatial domains generated using single-modal features: (C) isolated H&E image feature and (D) isolated spatial transcriptomic feature. (E) Quantitative assessment of clustering performance of the single-modal features versus the full multimodal HESTIA feature. (F) High-resolution subdomains generated by HESTIA using a refined parameter (K-means, k=9). (G) Volcano plot highlighting DEGs between the intra-tumoral domain 2 and domain 0. Genes exhibiting significant differential expression (FDR < 1e-4 and log2 fold change = 1.5) are marked in red. (H) Gene Ontology (GO) biological process pathway enrichment analysis. The bar chart displays the top 10 significantly enriched terms for both upregulated (red) and downregulated (blue) gene sets in domain 2 compared to domain 0. (I) Matrix plot illustrating the scaled mean expression profiles of selected signature DEGs across all spatial domains. (J) Spatial gene expression mapping of the top upregulated genes identified in domain 2 (*CCL19*, *IGLC2*, *IGLC3*, and *MZB1*).

### HESTIA dissects clinically relevant intratumoral heterogeneity in Visium HD colorectal cancer tissues

Having demonstrated HESTIA’s robust performance on high-resolution Stereo-seq data, we next evaluated its utility on another state-of-the-art spatial transcriptomics platform. We did so by applying HESTIA to two Visium HD colorectal cancer (CRC) samples, denoted as P1 and P2 (Figure 5A, J). Analyzed at a resolution of 8 μm, each sample contains over 500,000 bins. We used the reference annotations from the original study^11^ as a baseline for our evaluation. These annotations defined spatial cellular structures through unsupervised clustering and manual cell type labeling (Figure 5B, K). By integrating spatially resolved gene expression with paired H&E image, HESTIA successfully partitioned the bins into spatial domains representing the major cell types and structures inherent to the CRC microenvironment, including tumor, smooth muscle, fibroblasts, normal intestinal epithelial cells, and T cell-enriched zones (Figure 5C, L). When comparing HESTIA’s spatial domains against the reference labels, we observed that HESTIA further subdivided the broadly annotated tumor regions into multiple distinct subdomains. Consistent with their overarching identity, these subdomains exhibited uniformly high expression of *CEACAM6* (Figure 5D, M), an established CRC marker gene associated with poor overall survival and disease-free survival^32,33^. To understand the biological drivers behind HESTIA’s segmentation of these tumor regions, we investigated the molecular differences between the subdomains.

In sample P1, DEG analysis revealed distinct transcriptional signatures between the two primary tumor subdomains. The volcano plot (Figure 5E) and matrix plot (Figure 5F) showed that one specific subdomain (domain 4) had notably high expression of *REG1A*, *REG1B*, and *REG3A*. The REG gene family is tightly associated with CRC invasion, enhanced cellular proliferation, poor cancer differentiation, and apoptosis inhibition^34–36^. In contrast to the uniform mapping of *CEACAM6*, the tumor zone located in the center of the tissue section (domain 8) displayed a significantly lower expression of these REG family genes (Figure 5G–I).

Conversely, sample P2 did not exhibit the same differential expression of REG family genes in its HESTIA-identified tumor subdomains (Supplementary Figure 10). Instead, the segmentation in P2 appeared to be driven by microenvironmental cellular interactions. Specifically, one of the tumor subdomains (domain 8) was co-localized with *SPP1*+ macrophages (Figure 5N). *SPP1*+ macrophages are a specialized, tumor-specific myeloid population whose presence is strongly associated with immune exclusion, tumor progression, and worse clinical outcomes in CRC^37,38^.

**Figure 5.**
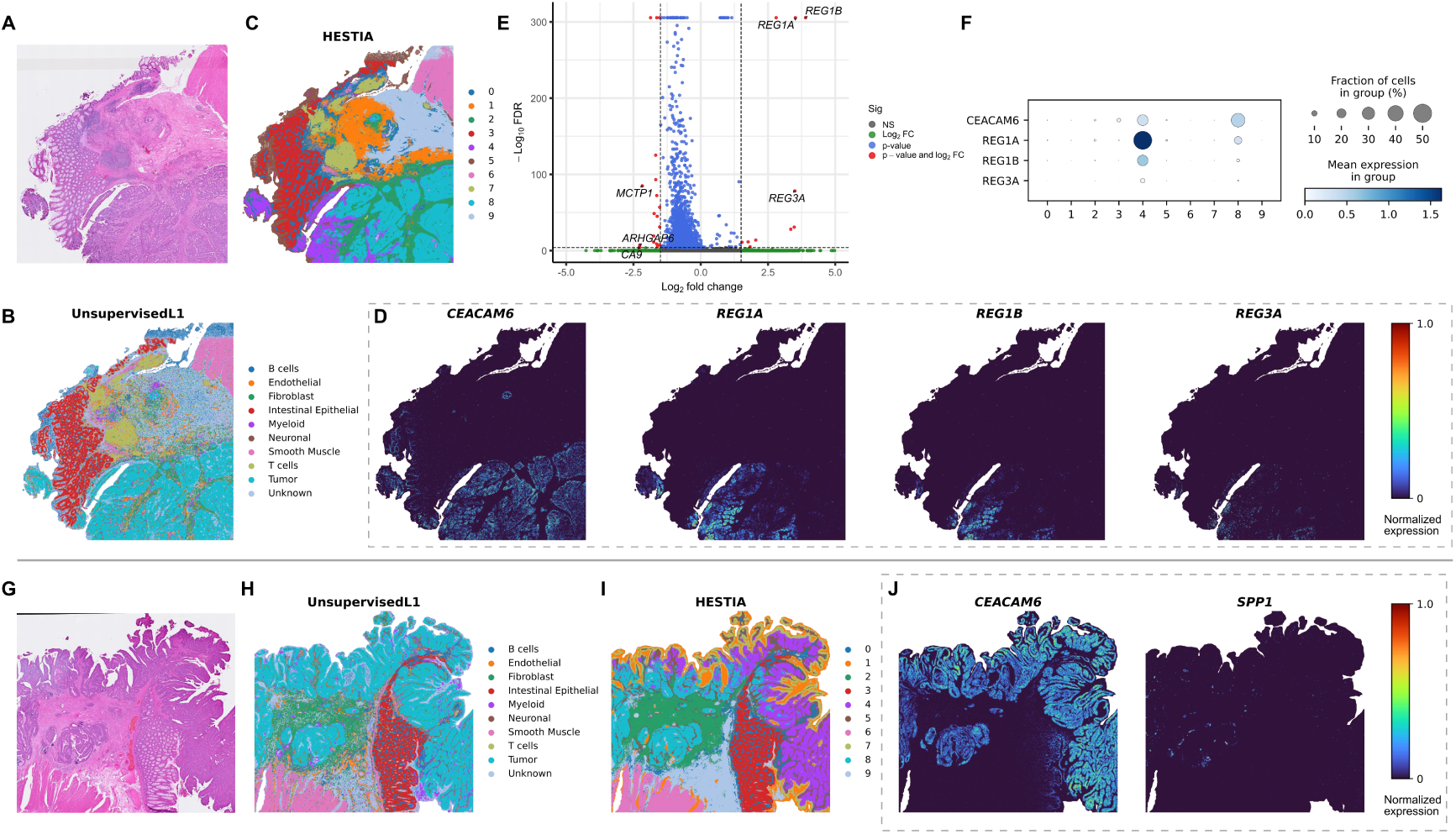
HESTIA dissects clinically relevant intratumoral heterogeneity in Visium HD colorectal cancer tissues. (A) H&E-stained histology image of human CRC sample P1. (B) Reference clustering and cell-type annotation results of sample P1 derived from the original dataset publication. (C) Spatial domains of sample P1 identified by HESTIA. (D) Spatial gene expression plots of the CRC marker gene *CEACAM6* and REG family genes (*REG1A*, *REG1B*, and *REG3A*) in sample P1. (E) Volcano plot illustrating DEGs between two primary intra-tumoral subdomains (domain 4 vs. domain 8) in sample P1. Genes exhibiting significant differential expression (FDR < 1e-4 and log2 fold change = 1.5) are marked in red. (F) Dot plot summarizing the expression profiles of *CEACAM6* and REG family genes across HESTIA-identified domains. (G) H&E-stained histology image of human CRC sample P2. (H) Reference clustering and cell-type annotation results of sample P2 derived from the original dataset publication. (I) Spatial domains of sample P2 identified by HESTIA. (J) Spatial gene expression plots of the CRC marker gene *CEACAM6* and the tumor-specific macrophage marker *SPP1* in sample P2.

### Ablation study and robustness analysis

HESTIA utilizes a dual-autoencoder system to process both the high-resolution and the aggregated low-resolution spatial transcriptomic data simultaneously. To evaluate the effectiveness of this architectural design, we conducted an ablation study with five configurations: a “highres-only” model that removes the low-resolution autoencoder entirely, an “identical-res” setup that replaces the low-resolution input with the exact high-resolution data, a “spatial-smoothing” baseline that replaces the low-resolution autoencoder with direct local averaging of the high-resolution expression matrix before the autoencoder, a “PCA” baseline that combines PCA of the expression matrix with PCA of the image features and directly applies clustering, and the full HESTIA model.

Using the previously established Stereo-seq mouse brain dataset, we compared the spatial domain identification performance of these configurations. The full HESTIA model achieved higher ARI and NMI scores compared to the ablated models and the baselines. For ASW and CHAOS, the PCA baseline performed slightly worse while the other configurations showed comparable results. For PAS, the spatial-smoothing baseline performed slightly worse and the PCA baseline performed substantially worse, whereas the full HESTIA model, the highres-only, and the identical-res configurations all maintained very low PAS values (Figure 6A). These results demonstrate that integrating parallel high- and low-resolution data improves spatial domain identification accuracy while preserving high spatial continuity.

To investigate whether the benefits of the dual-autoencoder system depend on data quality, we performed further ablation studies on a downsampled Stereo-seq mouse brain bin16 dataset with varying sequencing depths (see Methods). Using the mean ARI across the seven previously defined subregions as our evaluation metric, we assessed the clustering performance of the final multimodal joint feature (HESTIA), the isolated expression feature, and the isolated image feature independently. Since the image feature is unaffected by varying sequencing depths or ablation configurations, our analysis focused on the expression feature and the final joint feature.

When comparing the full HESTIA model against the highres-only setup, we found substantial gains in the clustering results of the isolated expression feature (Figure 6B). Notably, these performance gains showed a decreasing trend as sequencing depth increased (Figure 6C). Similar patterns were observed when comparing the full model against the identical-res configuration (Figure 6D, E). These findings indicate that the inclusion of both high- and low-resolution data has a greater impact on lower-quality data, suggesting that the dual-autoencoder system effectively stabilizes sparse molecular signals and mitigates the impact of gene dropout. As expected, while this robust transcriptomic stabilization greatly improves the expression-only representation, its effect on the final multimodal feature is more moderate, as the joint representation already benefits from the unchanged morphological feature provided by the histology image.

We further evaluated the robustness of HESTIA to key hyperparameters. Clustering accuracy slightly decreases as the bin size ratio increases (Supplementary Figure 11A), which is expected because a larger binning ratio substantially reduces the resolution of the low-resolution branch and may blur fine tissue structures. For the cross-resolution alignment weight α, performance remains stable across a wide range of values above the default (α = 1.0) but decreases when α is reduced below 1.0 (Supplementary Figure 11B), indicating that the cross-resolution alignment loss contributes meaningfully to the overall performance.

**Figure 6.**
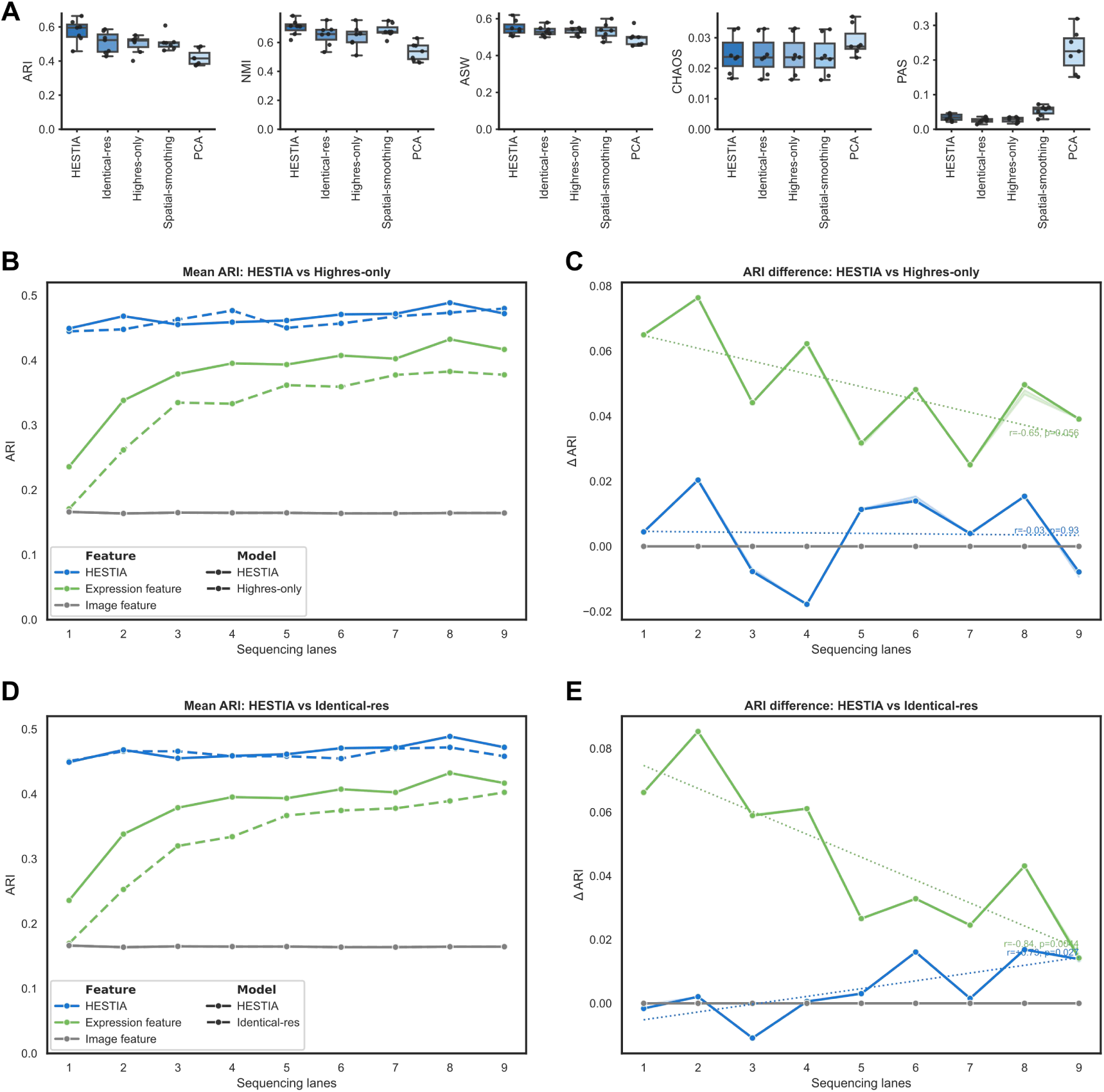
Ablation study. (A) Quantitative comparison of spatial domain identification performance across five ablation configurations: the complete multi-modal HESTIA model; an “Identical-res” model (where the aggregated low-resolution input is replaced by the exact high-resolution data); a “Highres-only” model (where the low-resolution autoencoder is entirely removed); a “Spatial-smoothing” (where the low-resolution autoencoder is removed, and the high-resolution data are smoothed by replacing each spot with its local average); and a “PCA” (which clusters on the PCA of the expression matrix combined with PCA of the HIPT features, with all autoencoders removed). Boxplots display clustering accuracy (ARI, NMI) and spatial continuity (ASW, CHAOS, PAS) for HESTIA and the compared algorithms. Higher ARI, NMI, and ASW scores denote superior performance, whereas lower CHAOS and PAS values indicate greater spatial coherence. The center lines of the boxplots represent the median, the box boundaries represent the IQR, and the whiskers extend to 1.5× IQR. (B) Mean ARI comparing the full HESTIA framework (solid lines) against the “Highres-only” ablated model (dashed lines) across varying sequencing depths. The performance of the final integrated multimodal joint feature (blue), the isolated expression feature (green), and the isolated image feature (grey) was evaluated. (C) The performance gain of the full HESTIA model compared to the “Highres-only” model across varying sequencing depths. Dotted lines represent linear regression trends. Pearson correlation coefficients (r) and associated p-values (Wald test with t-distribution) are shown. (D) Mean ARI comparing the full HESTIA framework (solid lines) against the “Identical-res” ablated model (dashed lines) across varying sequencing depths. (E) The performance gain of the full HESTIA model compared to the “Identical-res” model across varying sequencing depths.

### HIPT feature generalizability across tissue types

The HViT model we used in HESTIA is Hierarchical Image Pyramid Transformer (HIPT, see Methods for details). HIPT was originally pretrained on human pathology images from TCGA, and its application to mouse brain H&E involves a cross-species domain shift. To evaluate whether this shift affects feature quality, we compared the HIPT embeddings of the mouse brain sample with those of the human cancer samples (lung adenosquamous carcinoma and colorectal cancer) used in this study. In the token-level PCA projection, mouse and human histology tokens showed substantial overlap rather than forming two sharply separated clusters, indicating that the domain difference does not dominate the representation space (Supplementary Figure 12A). When colored by sample identity, the mouse brain tokens occupied a coherent region while remaining embedded within the broader distribution of human pathology samples, rather than appearing as a fully isolated cluster (Supplementary Figure 12B). Consistently, sample-level cosine similarity analysis showed that the mouse brain sample retained high similarity to the human samples, with cosine similarities ranging from 0.88 to 0.94 (Supplementary Figure 12C). Although the mouse sample was somewhat less similar to some human samples than the human samples were to one another, its similarity remained high overall. Quantitatively, the silhouette score in the PCA space was approximately zero (−0.002), confirming the absence of systematic domain separation between mouse and human tokens. These results suggest that the pretrained HIPT features generalize reasonably well to mouse brain H&E, and that the domain shift does not lead to a complete failure of the representation. Detailed methods for this analysis are provided in the Supplementary Methods.

### Adapting HESTIA to alternative histology feature extractors

To evaluate the flexibility of HESTIA with respect to the underlying histology feature extractor, we substituted HIPT with four histology foundation models: Prov-GigaPath^39^, Virchow2^40,41^, UNI and UNI2-h^42^. These models were pretrained on hundreds of thousands to millions of whole-slide images (WSIs) and contain approximately 300 million to 1 billion parameters, both representing one to two orders of magnitude greater scale than HIPT. None of these models adopts a hierarchical two-stage architecture comparable to that of HIPT. Additionally, UNI2-h and Virchow2 use a different image token size (14 × 14 pixels) compared to HIPT, UNI, and Prov-GigaPath (16 × 16 pixels), introducing further architectural variation. For each alternative model, we applied the same default hyperparameters originally tuned for HIPT, without any model-specific re-optimization. Across the seven mouse brain subregions, these variants continued to outperform the other multimodal algorithms evaluated in our benchmark, despite not reaching the performance of the original HIPT configuration (Supplementary Figure 13). These findings indicate that the core design of HESTIA, encompassing the dual-autoencoder framework and the modality fusion strategy, remains effective when paired with different histology feature extractors. However, fully exploiting the capacity of a more powerful foundation model is likely to require dedicated hyperparameter optimization.

## Discussion

The transition of spatial omics toward high-throughput, subcellular resolution provides unprecedented opportunities for dissecting complex tissue architectures. As datasets scale to include multi-million cells, computational bottlenecks and inherent transcriptomic sparsity severely challenge existing analytical toolkits. Although recent methods such as BINARY and MENDER achieve excellent scalability, they rely exclusively on a single transcriptomic modality. Today, many modern spatial omics platforms routinely co-register tissue with paired H&E images. Integrating gene expression with morphological features enables superior tissue characterization and opens new avenues for clinical and diagnostic applications^43,44^. To bridge this gap, we developed HESTIA, a scalable multimodal framework that integrates matched histology imagery with spatial gene expression for robust spatial domain identification.

HESTIA is fundamentally designed to tackle the challenges posed by high-resolution spatial transcriptomic data: feature sparsity and extreme scale. It accomplishes this through two specific architectural designs. First, HESTIA employs a novel dual-autoencoder system to alleviate severe gene dropout. As demonstrated by our ablation study, this cross-resolution consistency primarily stabilizes the expression representation under low-depth or sparse conditions. While its contribution to the final multimodal feature is modest (since the image modality is unaffected by sequencing depth and already provides a stable complementary signal), it still offers meaningful improvements when transcriptomic data quality is limited. Notably, the robustness of the multimodal feature to varying sequencing depth, driven by the stable histology signal, further reinforces the value of integrating complementary modalities. Second, HESTIA achieves extraordinary computational efficiency by intentionally avoiding memory-intensive deep learning schemes. Unlike existing multimodal algorithms that explicitly build memory-demanding graph neural networks based on spot proximity, HESTIA indirectly encodes spatial context via its hierarchical image model and dual-resolution constraint. While this design choice sacrifices the explicit local inductive bias inherent to spatial graphs, it fundamentally circumvents the quadratic memory bottlenecks that prevent competing methods from scaling to massive datasets. This trade-off is deliberate: by omitting graph construction, HESTIA can process datasets such as a two-million-bin human lung carcinoma slice on a single standard GPU, whereas other multimodal algorithms fail due to out-of-memory errors.

This optimized design enables the high-fidelity mapping of complex tissue structures. In the mouse brain dataset, HESTIA successfully delineated complex anatomical structures at both the high-resolution grid level (bin20) and the single-cell level (cellbin). It resolved subtle structural margins, such as the CA1/CA3 stratum pyramidale and the dentate gyrus, significantly outperforming other models that blurred these boundaries. In clinical oncology settings, HESTIA provided profound diagnostic insights. In colorectal cancer samples, HESTIA captured intra-tumoral heterogeneity, identifying an aggressive tumor boundary enriched with *REG* family genes and distinguishing a distinct tumor state structured by its proximity to pro-tumorigenic *SPP1*+ macrophages. Additionally, in a massive lung adenosquamous carcinoma dataset, HESTIA identified an immunologically active niche driven by B-cell immunity networks within the squamous cell region. We note that the ARI of 0.33 and NMI of 0.35 on this sample are modest compared to those in the mouse brain. However, these values are consistent with published benchmarks: on the same lung sample, StereoMM achieved a median ARI of 0.32 and NMI of 0.34 (albeit at bin100 resolution and in four subregions)^10^. Similarly, studies on specific breast cancer^45–47^, prostate cancer^48^, and lymph node^49^ datasets have reported best ARI values of 0.22 to 0.35, and another study on colorectal cancer^50^ reported that the agreement between different methods ranged from 0.32 to 0.6. These results indicate that the inherent complexity of heterogeneous pathological tissues often yields modest agreement metrics, and HESTIA’s performance is comparable to the state-of-the-art.

Despite its outstanding performance, HESTIA has several limitations. First, our spatial aggregation strategy groups adjacent bins or cells into simple squares to construct the low-resolution input. While this approach is computationally efficient, the rigid grid inherently ignores the boundaries of distinct cell types, potentially mixing heterogeneous adjacent cells into a single bin. Although the current impact is minimal since large bins contain only four high-resolution sub-bins, incorporating cell-type-aware grouping strategies could further refine spatial domain boundaries^51^, provided the additional computational overhead is properly managed. Second, while HESTIA has demonstrated exceptional performance on sequencing-based spatial transcriptomics platforms, its application to image-based platforms requires further evaluation. Because image-based technologies typically capture fewer gene types but have significantly higher sensitivity^52^, our dual-autoencoder parameters may require specific tuning to adapt to these denser transcriptomic profiles. Third, our quantitative benchmarks with ground truth annotations are limited to mouse brain. Extending the evaluation to other tissue types will be important as suitable datasets become available. Fourth, HESTIA’s hyperparameters are optimized specifically for the HIPT feature extractor. When an alternative foundation model is substituted without re-optimizing these hyperparameters, the resulting performance is suboptimal, although the framework still outperforms competing multimodal algorithms. Finally, as a multimodal algorithm that relies on paired H&E imagery, HESTIA is sensitive to the optical quality of the input. Histological artifacts such as tissue folds, tears, or inconsistent staining can introduce morphological noise that perturbs the HViT embeddings, potentially leading to spatial domain misclassifications in affected regions.

## Methods

### Stereo-seq mouse brain data acquisition

A 6-week-old C57BL/6 mouse was used to collect brain tissue. The tissue sample was frozen and embedded in OCT. Then, cryosections were cut at a thickness of 10 μm using a Leica CM1950 cryostat. The sample preparation and sectioning procedures followed the Guide for Fresh Frozen Samples on Stereo-seq Chip Slides (Document No. STUM-SP001). Subsequent steps, including hematoxylin and eosin (H&E) staining, in situ reverse transcription, amplification, library construction, and sequencing, followed the User Manual of Stereo-seq Transcriptomics Set for Chip-on-a-Slide V1.3 (STOmics, #211ST13114). Briefly, the tissue section was adhered to a Stereo-seq chip (generated by BGI, China) and fixed in precooled methanol. Prior to tissue permeabilization, H&E staining was performed. The RNA was then released from the permeabilized tissue, captured by the DNB, and underwent in situ reverse transcription. After reverse transcription, the tissue section was removed to release the cDNA and amplify it. The cDNA was purified using VAHTS^TM^ DNA Clean Beads (Vazyme, #N411-02), strictly following the User Manual of Stereo-seq 16 Barcode Library Preparation Kit (STOmics, #101KL11160). For library construction, 100 ng of the cDNA was used for fragmentation and amplification. The PCR products were then purified using VAHTS^TM^ DNA Clean Beads (Vazyme, #N411-02). The purified PCR products were then used for DNB production, and the libraries were sequenced on the MGI sequencer.

The raw sequencing data were processed using the SAW pipeline. Low-quality reads were filtered and the CID sequences were mapped to the chip coordinates. Then, the reads were aligned to the mm10 genome. To eliminate PCR duplicates, UMIs for the same CID and gene were collapsed. Spatial gene expression matrix was then generated by integrating gene counts with coordinates.

### Data preprocessing

We used Scanpy^53^ to preprocess the spatial transcriptomics data. Specifically, genes expressed in fewer than three cells were filtered out. Then the counts were library-size normalized and log-transformed, followed by scaling to unit variance. PCA was applied and the top 64 principal components were extracted, yielding compact expression representation for downstream modeling.

The H&E image must be aligned with the spatial transcriptomics data so that each pixel in the image corresponds to a DNB (or bin1) in the Stereo-seq data or a 0.5 μm x 0.5 μm pseudo-square in the Visium HD data. For Stereo-seq data, we used the SAW software to align the H&E image and perform cell segmentation. The output GEF format files were then converted to anndata format using Stereopy^54^. For Visium HD data, we first used SpatialData^55^ to convert it to anndata format. Then the image and the spatial transcriptomic data were aligned with the square grid using similarity transformation (i.e., rotation, shifting, and uniform scaling).

### Multiscale histology feature extraction

We used a Hierarchical Vision Transformer (HViT) to derive histology embeddings that retain both local structure and global tissue context without patch-by-patch tiling. To leverage the large-scale publicly available histology datasets, we adopted a pretrained model named Hierarchical Image Pyramid Transformer (HIPT), which was pretrained on more than 10,000 gigapixel WSIs from The Cancer Genome Atlas (TCGA)^56^. This HViT model extracts features in two hierarchical stages, moving from local cell-level patterns to broader tissue context. First, it performs patch tokenization by partitioning the WSI into patches of 256×256 pixels and decomposing each patch into 16×16 non-overlapping tokens. Second, a local Vision Transformer (ViT) extracts a CLS summary embedding that captures fine-grained cellular and structural attributes from each patch. These patch-level CLS embeddings are then organized within 4096×4096 pixel regions and passed as sequences to a global ViT. The global ViT models long-range spatial dependencies and inter-patch interactions, thereby contextualizing local features within broader tissue structure and yielding region-informed embeddings. We extract the embedding for each 16x16 pixel token and concatenate it with its corresponding CLS embedding to create a feature for each 16x16 pixel image patch.

To compress and regularize the image embeddings, we train a lightweight autoencoder on the HViT features. The loss function for the autoencoder is expressed as follows:

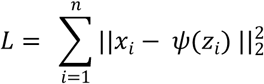

where *x_i_* is the input image feature, *z_i_* is the latent feature from the bottleneck layer, and *ψ* is the decoder. Once training is complete, the decoder is discarded and the latent feature is used as the representation of the image patch for modality fusion.

### Multiscale spatial transcriptomics representation learning

To exploit complementary information across spatial resolutions and to alleviate information loss due to gene capture sparsity at high resolution, we construct paired high- and low-resolution views of the spatial transcriptomic data. For the high-resolution view, we obtain the raw expression data from the input spatial transcriptomics, followed by library-size normalization, log transformation and scaling to unit variance. Then we extract the top 64 principal components from the data to use as expression features. For the low-resolution view, we first aggregate the high-resolution raw expression data into coarser bins with a specific bin size ratio while preserving coordinates. The default bin size ratio is 2, which means a low-resolution bin consists of 2x2 high-resolution bins. For cellbin data, we aggregate cells within a 40 x 40 square into a bin40 data, assuming each cell approximates the size of a bin20. We then preprocess the low-resolution data using the same procedures as the high-resolution data.

Next, we construct a dual-autoencoder system in which separate autoencoders are trained for high- and low-resolution features. In addition to the reconstruction loss for each autoencoder, we introduce a cross-resolution alignment loss to minimize discrepancies between the high- and low-resolution latent representations at matched locations. Specifically, the reconstruction losses are:

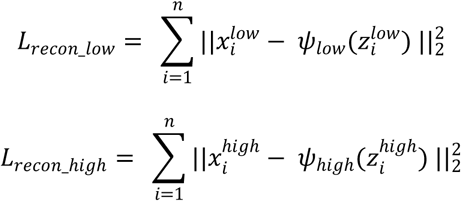

where 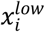 is the expression feature of the *i* th low-resolution bin, 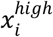 is the expression features of the *i*th low-resolution bin, which consists of approximately 2x2 high-resolution bins (*i.e.,* for cellbin data and bins at tissue border, the number may vary). ψ*_low_* and ψ*_high_* are the decoders, 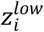 and 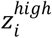 are the latent representations for each resolution. The cross-resolution alignment loss is:

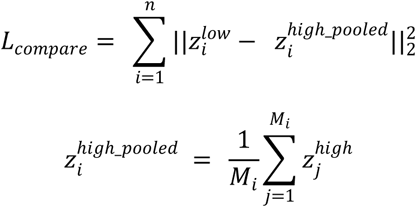

where 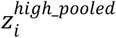 is the high-resolution latent representation averaged by the number of high-resolution bins in a low-resolution bin (*M_i_*).

The total loss of the dual-autoencoder system is a weighted sum of the reconstruction losses and the cross-resolution alignment loss:

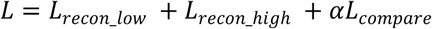

where *a* is a hyperparameter that determines the weight given to the cross-resolution alignment loss relative to the reconstruction loss, and is set to 1 by default.

Both the image autoencoder and the transcriptomics dual-autoencoder system are trained using the Adam optimizer^57^ with a weight decay of 0.0001, and a cosine annealing learning rate schedule initiated by a learning rate of 0.0005.

### Modality fusion

Following modality-specific encoding, we integrate the transcriptomic and histology representations using computationally efficient fusion operators, similar to those in MISO^19^. First, the Kronecker product of the image latent representation and the high resolution transcriptomic latent representation *z^high^* are computed, giving the interaction feature matrix:

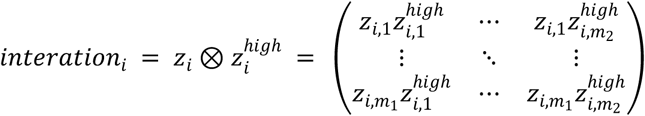

where *m_1_* is the dimension of the image latent representation, *m_2_* is the dimension of the high-resolution transcriptomic latent representation. The interaction feature matrix is then flattened for all spots, followed by dimension reduction via PCA. Next, the two representations are standardized to a mean of 0 and a variance of 1, and then concatenated to form a single representation:

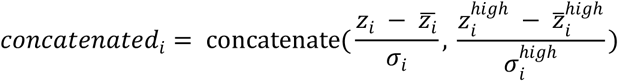

where 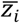 is the mean value of image latent representation *z_i_*, *σ_i_* is the standard deviation value of *z_i_*; 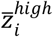 is the mean value of high resolution transcriptomic latent representation 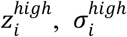 is the standard deviation value of 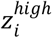.

Finally, the dimension reduced interaction term is concatenated with the two standardized representations:

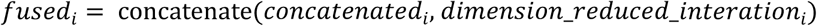

By default, this final fused representation is used for downstream tasks such as unsupervised clustering, but we also provide options to use features from a single modality, or the interaction term or the concatenation term alone. This offers users tailored control based on their specific research needs.

### Benchmarking of different multimodal spatial domain identification algorithms

We compared the performance of HESTIA with eight other multimodal algorithms. To make a fair comparison, we clustered the learned representations from all algorithms using K-means and ignored all post-processing steps (e.g., cluster refinement in ConGI and SpaGCN). This ensures that the comparison focuses on the core mechanism of the algorithms instead of the differences in clustering methods. All algorithms were run on a Linux server with 56 CPU cores, 2000 GB of memory, and an NVIDIA T4 GPU with 16 GB of GPU RAM.

### Differentially expressed genes (DEGs) and enrichment analysis

Differentially expressed genes were computed using the Wilcoxon rank-sum test implemented in Scanpy. The p-values were then adjusted by Benjamini-Hochberg procedure^58^. Both pathway enrichment of the DEGs and GSEA analysis were performed against the Gene Ontology (GO)^59^ biological process database using the gseapy^60^ implementation.

### Downsampling of Stereo-seq data

We used mouse brain data that were downsampled by different sequencing lanes for ablation study. The full dataset was acquired from nine lanes and contained a total of 2.6 billion sequenced reads. Downsampling results in spatial transcriptomics data of varying quality. The median gene counts for each downsampled bin16 data are shown in Supplementary Table 2.

## Key Points

- HESTIA is a multimodal computational framework that jointly analyzes high-resolution spatial transcriptomics and matched histology images through a dual-autoencoder system guided by cross-resolution consistency constraints and a hierarchical vision transformer.
- HESTIA demonstrates robust analytical performance by overcoming the severe memory constraints and transcriptomic sparsity of subcellular-resolution platforms, consistently achieving superior clustering accuracy and spatial continuity compared to eight competing algorithms while seamlessly processing full-slice datasets exceeding two million spots on a single GPU with 24 GB of memory.
- Applied to mouse brain and human cancer tissues, HESTIA resolves fine anatomical structures at single-cell resolution and uncovers clinically relevant intratumoral heterogeneity, including B-cell-enriched immune niches in lung cancer and *SPP1⁺* macrophage-driven tumor microenvironments in colorectal cancer.

## Acknowledgements

We thank Bingying Luo, Fei Teng and Jiajun Zhang for their assistance in data preparation. We also thank the China National GeneBank for providing data storage support for this study.

## Author contributions

Z.Z. and A.C. conceived the study. Z.Z. implemented the algorithm and ran all the experiments. Z.Z. and X.Z. analyzed the results of the manuscript. J.G. and S.L. provided the mouse brain data. X.Z. annotated the domains for the mouse brain data. S.L. and A.C. supervised the study. All authors contributed to writing and reviewing the manuscript.

## Data availability

The Stereo-seq mouse brain data have been deposited into CNSA with accession number CNP0009334 (https://db.cngb.org/data_resources/project/CNP0009334/). Stereo-seq human lung adenosquamous carcinoma data can be accessed from GSA-human (https://ngdc.cncb.ac.cn/gsa-human/) under Project HRA004240. 10x Visium HD human colorectal cancer data are available at https://www.10xgenomics.com/datasets/visium-hd-cytassist-gene-expression-libraries-of-human-crc.

## Code availability

The python implementation of HESTIA is available on GitHub at https://github.com/zzhongzz/HESTIA. The repository also includes the scripts for benchmarking performance and computational resources, as well as notebooks for preprocessing Visium HD data.

