## Supplementary for "HESTIA: Scalable Multimodal Integration of Histology and High-Resolution Spatial Transcriptomics for Robust Spatial Domain Identification"

### Supplementary Methods

#### Cell type annotation of mouse brain data

A MERFISH mouse brain section from the Allen Brain Cell Atlas[1] that was morphologically similar to our Stereo-seq sample was selected, and cell type proportions were estimated based on the accompanying annotation. Processed single-cell RNA-sequencing data (10x Genomics Chromium v3 assays) from the Allen Brain Cell Atlas[2] were obtained, and approximately 52,000 cells were randomly subsampled to match the estimated cell type proportions. These cells were used to construct reference signatures for cell2location[3]. Two reference signatures were built using the "subclass" and "supertype" label sets from the single-cell data and applied to annotate the cellbin level Stereo-seq data.

#### Cell type annotation of human lung adenosquamous carcinoma data

Processed single-cell RNA-sequencing data from Dong *et al.*[4] were obtained and used to construct reference signatures for cell2location. The "cell_type" label sets were used and applied to annotate the Stereo-seq human lung adenosquamous carcinoma data.

#### Clustering of human lung adenosquamous carcinoma data using alternative methods

The cell2location annotation score (q05_cell_abundance_w_sf) and the principal components (n_pcs=50) of the expression matrix were used for clustering. Three clustering algorithms—K-means, Leiden[5], and Louvain[6]—were applied using two parameter settings each: k=5 or k=9 for K-means, and resolution=0.3 or resolution=1 for Leiden and Louvain.

#### HIPT embedding space analysis

A qualitative embedding-space analysis was performed using cached raw HIPT features extracted from mouse brain and cancer histology images. For each sample, 10,000 spatial tokens were randomly selected from the cached HIPT feature maps to ensure balanced representation across samples of varying image sizes. The sampled token embeddings were combined across all samples and projected into two dimensions using principal component analysis (PCA). In addition, a sample-level representation was computed for each image by pooling token embeddings across the full image. Pairwise similarity between samples was quantified using cosine similarity. Domain separation in the PCA projection was quantified using the silhouette score, which measures the degree of cluster separation between mouse and human tokens.

### Supplementary Tables

**Supplementary Table 1 Characteristics of the Stereo-seq mouse brain dataset across varying spatial resolutions.**
The table provides the total number of bins and the median gene counts per bin for each evaluated bin size, ranging from 256 down to 16.

| Bin size | Number of bins | Median gene counts per bin |
|---|---|---|
| 256 | 3,414 | 10,826 |
| 128 | 13,261 | 6,578 |
| 64 | 52,058 | 2,981 |
| 32 | 205,971 | 1,057 |
| 16 | 818,578 | 312 |

**Supplementary Table 2 Median gene counts associated with varying sequencing depths.**
The table shows the median gene counts per bin corresponding to the number of sequencing lanes used, representing different sequencing depths for the down-sampled Stereo-seq mouse brain bin16 dataset.

| Number of lanes | Median gene counts per bin |
|---|---|
| 1 | 108 |
| 2 | 156 |
| 3 | 196 |
| 4 | 225 |
| 5 | 252 |
| 6 | 272 |
| 7 | 289 |
| 8 | 304 |
| 9 | 312 |

### Supplementary Figures

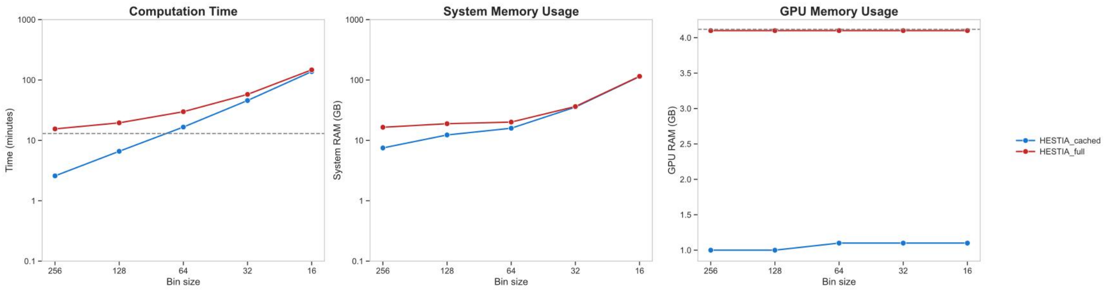


**Supplementary Figure 1 Computational resource evaluation of HESTIA with and without caching.**

Line plots compare the computation time, system memory usage, and GPU memory usage of the complete HESTIA pipeline (HESTIA_full) versus its modular caching mode (HESTIA_cached) across decreasing bin sizes from 256 to 16.

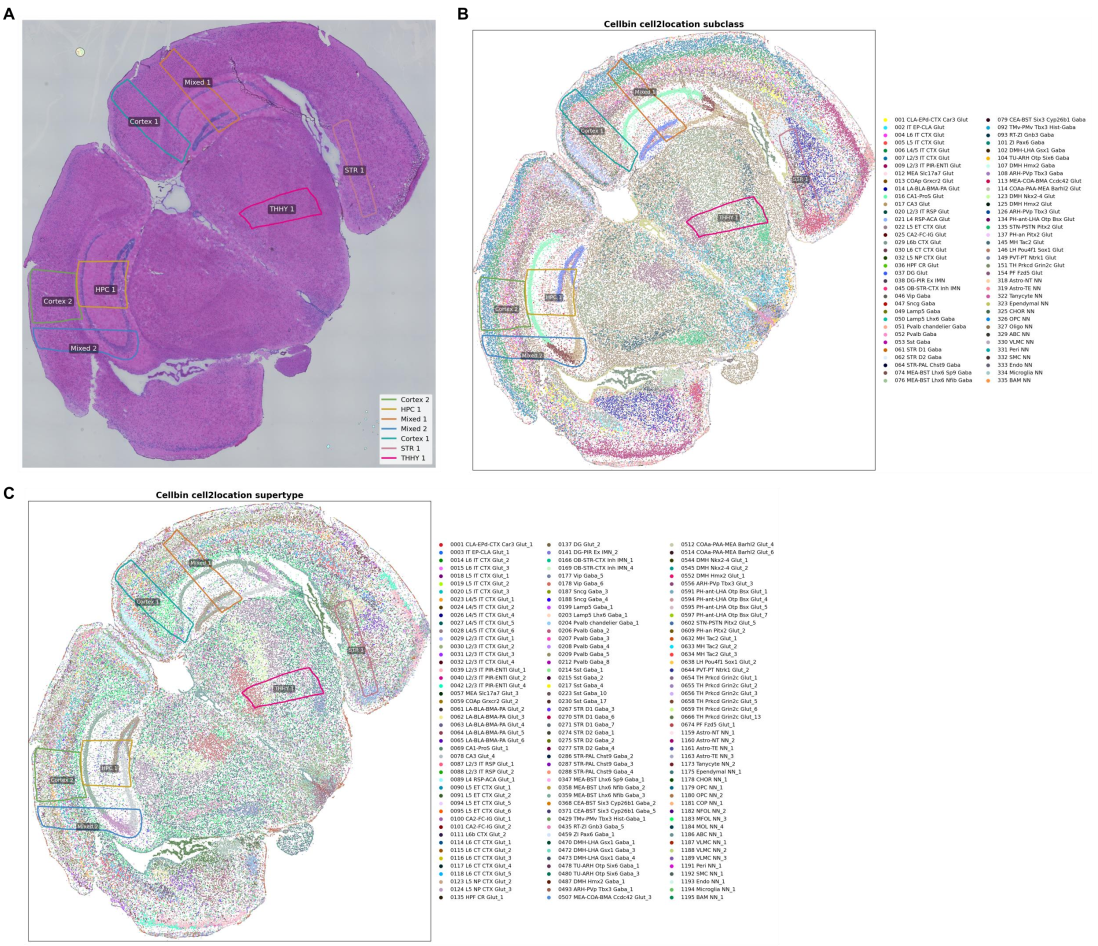


**Supplementary Figure 2 Locations of the seven selected subregions within the mouse brain dataset.**

(A) An H&E-stained whole-slice histology image of the mouse brain highlights the seven distinct subregions (Cortex 1, Cortex 2, HPC 1, Mixed 1, Mixed 2, STR 1 and THHY 1) encompassing the cortex, hippocampus, striatum, thalamus and hypothalamus that were utilized to evaluate spatial domain detection performance. (B) Cell type annotations generated by Cell2location, which used the "subclass" cell types. (C) Cell type annotations generated by Cell2location, which used the "supertype" cell types, showing a more refined classification.

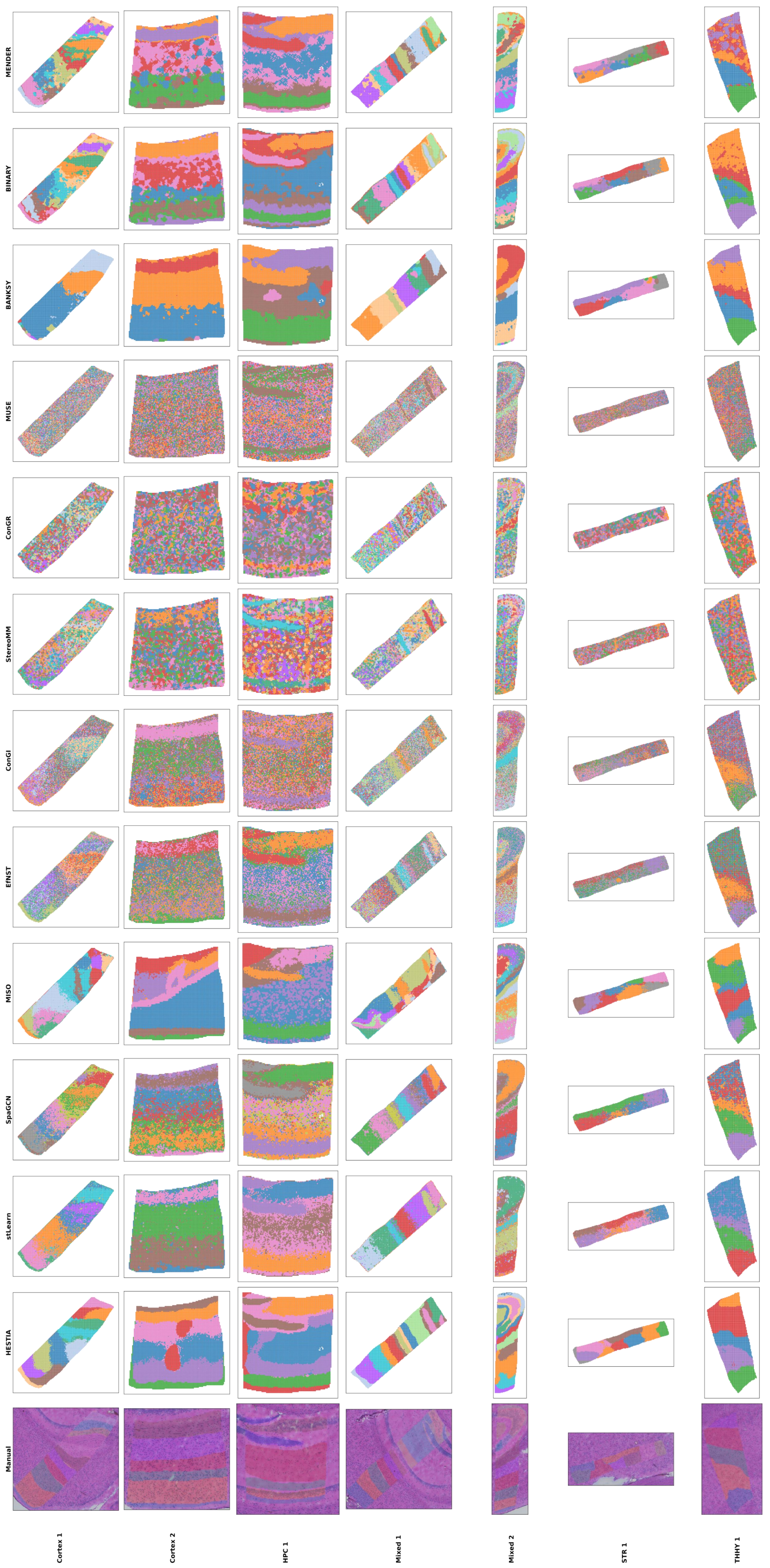


**Supplementary Figure 3 Spatial domain identification results for seven selected bin20 mouse brain subregions.**

Spatial scatter plots illustrate the domain clustering assignments generated by HESTIA and eight other multimodal algorithms, as well as three single modal algorithms, alongside manual reference annotations, across the Cortex 1, Cortex 2, HPC 1, Mixed 1, Mixed 2, STR 1 and THHY 1 subregions.

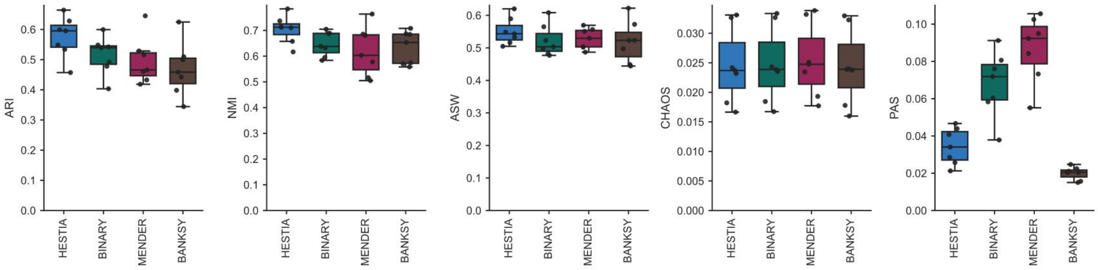


**Supplementary Figure 4 Quantitative evaluation of HESTIA and single-modal algorithms across seven mouse brain subregions at bin20 level.**

Boxplots display the clustering accuracy metrics (ARI and NMI) and the spatial continuity metrics (ASW, CHAOS, and PAS). Higher ARI, NMI, and ASW scores denote superior performance, whereas lower CHAOS and PAS values indicate greater spatial coherence. The center lines of the boxplots represent the median, the box boundaries represent the interquartile range (IQR), and the whiskers extend to 1.5× IQR.

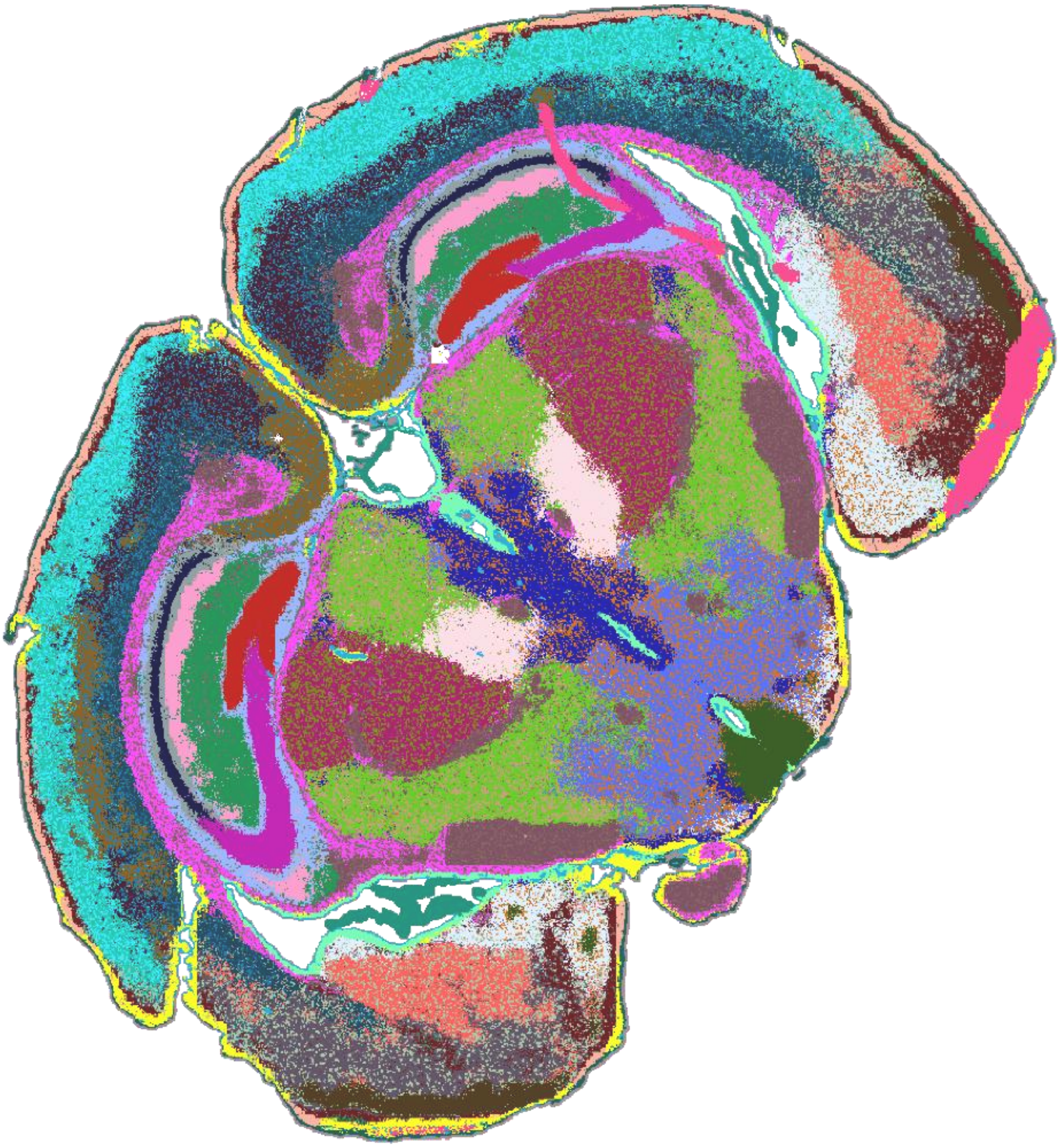

**Supplementary Figure 5 Whole-slice spatial domain identification on the bin20 mouse brain dataset.**

Spatial scatter plot visualizing the clustering assignments generated by HESTIA across the entire Stereo-seq mouse brain tissue section at the bin20 resolution.

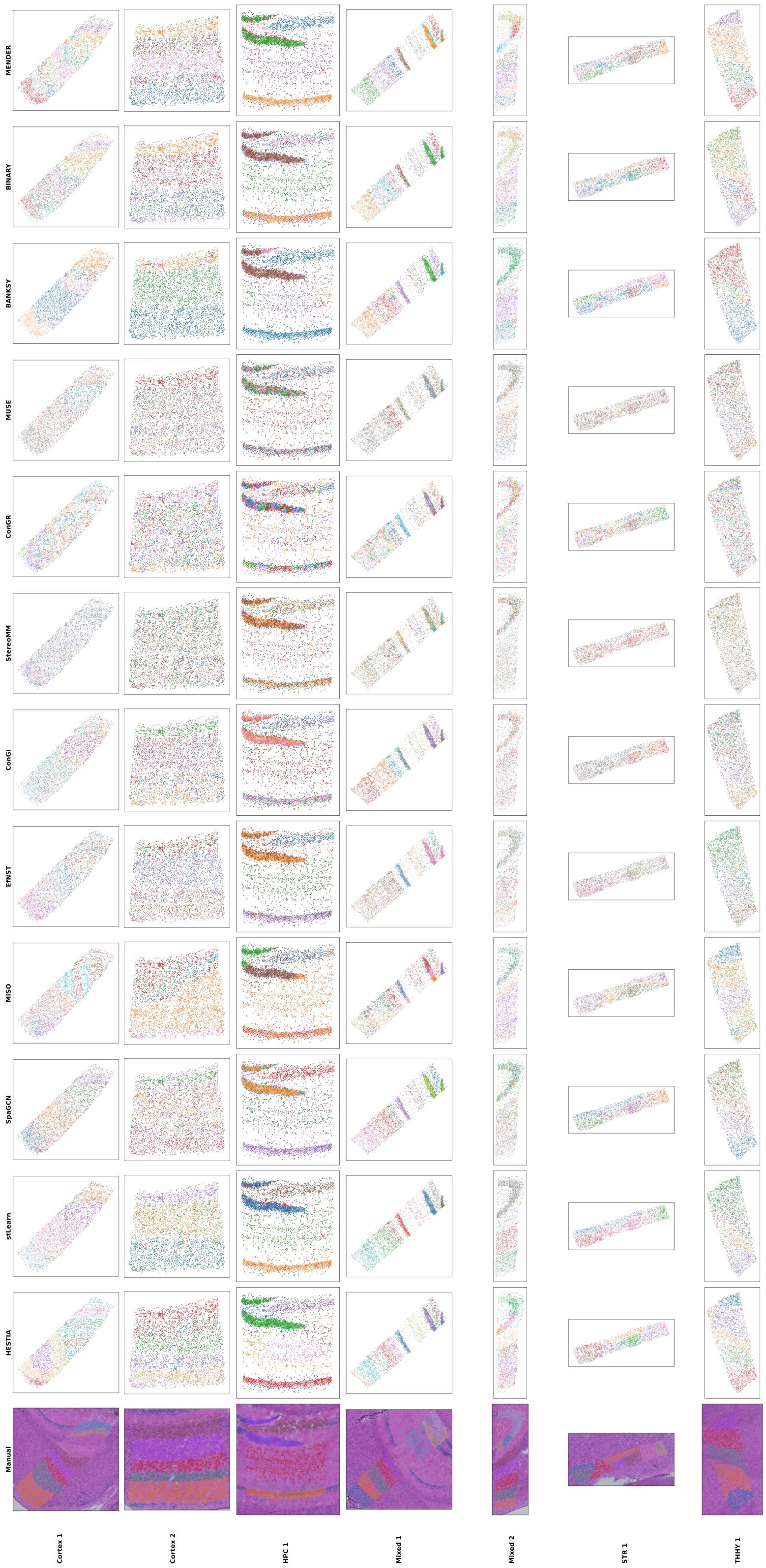


**Supplementary Figure 6 Spatial domain identification results for seven selected single-cell (cellbin) mouse brain subregions.**

Spatial scatter plots illustrate the domain clustering assignments generated by HESTIA and eight other multimodal algorithms, as well as three single modal algorithms, alongside manual reference annotations, across the Cortex 1, Cortex 2, HPC 1, Mixed 1, Mixed 2, STR 1 and THHY 1 subregions.

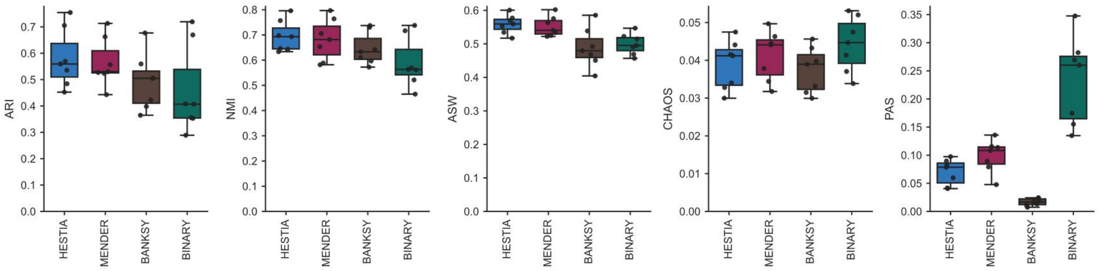


**Supplementary Figure 7 Quantitative evaluation of HESTIA and single-modal algorithms across seven mouse brain subregions at cellbin level.**

Boxplots display the clustering accuracy metrics (ARI and NMI) and the spatial continuity metrics (ASW, CHAOS, and PAS). Higher ARI, NMI, and ASW scores denote superior performance, whereas lower CHAOS and PAS values indicate greater spatial coherence. The center lines of the boxplots represent the median, the box boundaries represent the interquartile range (IQR), and the whiskers extend to 1.5× IQR.

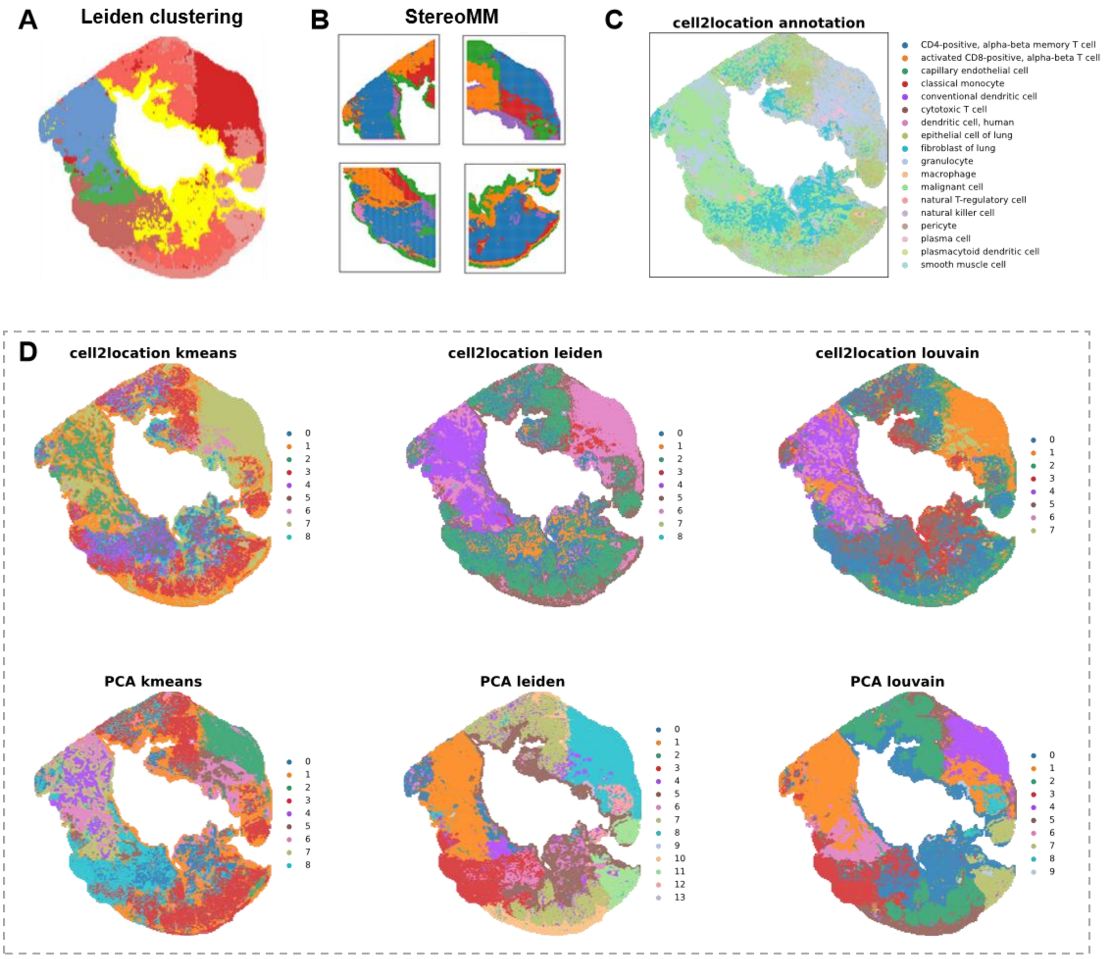


**Supplementary Figure 8 Spatial domain identification on the human lung adenosquamous carcinoma dataset using alternative methods.**

(A) Leiden clustering result obtained from the original study. (B) StereoMM clustering results obtained from the original study. It was performed on four partitioned subregions due to memory constraints. (C) Cell type annotations generated by Cell2location. (D) Clustering results based on the Cell2location annotation scores (first row) and the PCA of the expression matrix (second row).

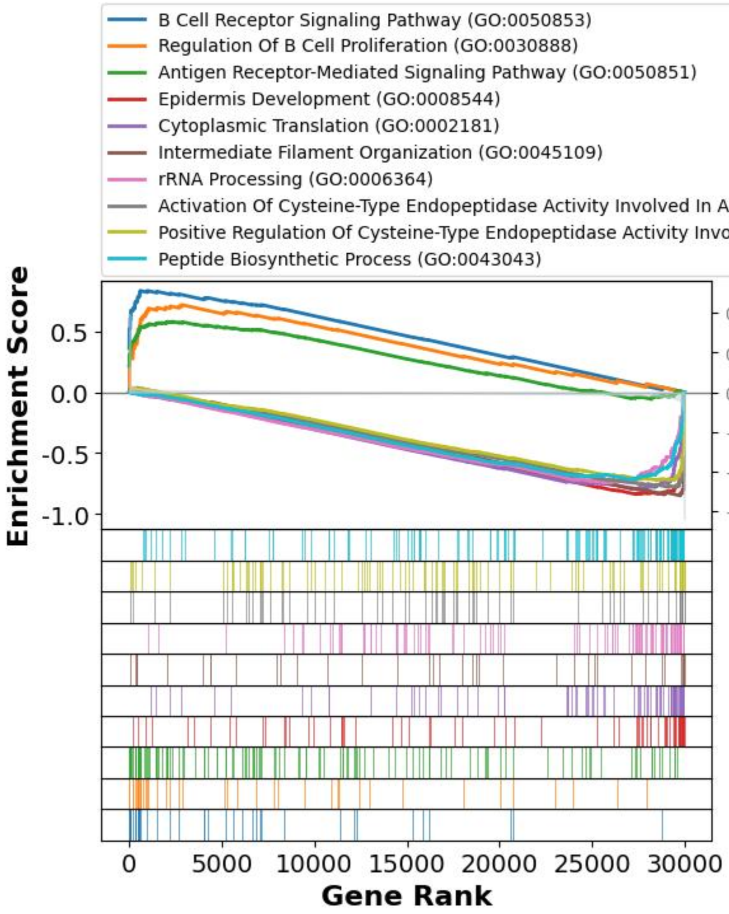


**Supplementary Figure 9 GSEA confirming significantly altered biological pathways between domain 2 and domain 0.**

Curves represent the running enrichment score for the top 10 enriched GO terms, while vertical tick marks denote the occurrence of genes from each GO term within the ranked input statistic list.

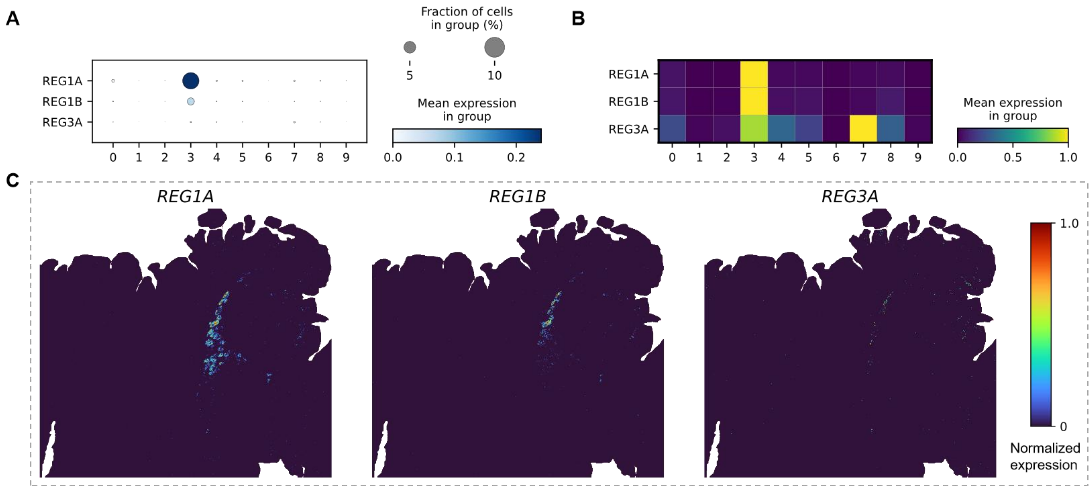


**Supplementary Figure 10 Expression profiles of REG family genes in the Visium HD human colorectal cancer sample P2.**

(A) Dot plot and (B) matrix plot showing the expression levels of *REG1A*, *REG1B*, and *REG3A* across the HESTIA-identified spatial domains. (C) Spatial gene expression plots of *REG1A*, *REG1B*, and *REG3A*.

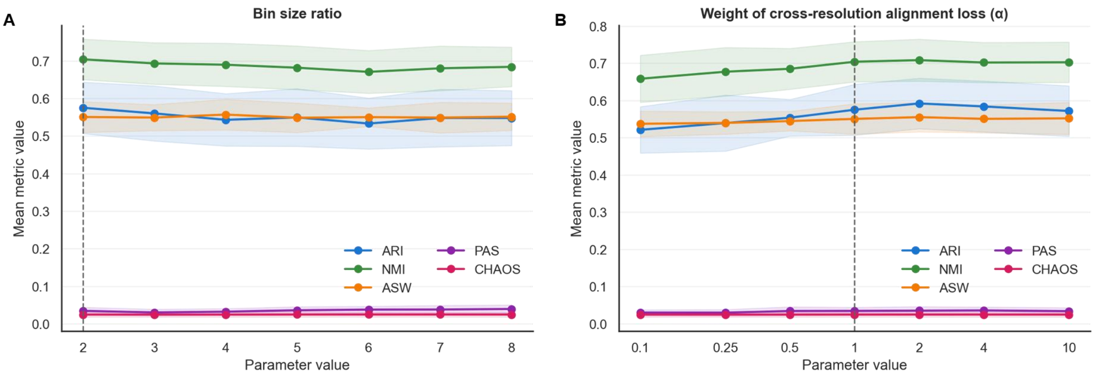


**Supplementary Figure 11 Hyperparameter robustness analysis of HESTIA.**

(A) Mean performance of HESTIA across seven samples under different bin size ratio settings. Curves show the mean values of ARI, NMI, ASW, PAS, and CHAOS, and shaded areas indicate one standard deviation across samples. The vertical dashed line marks the benchmark setting (bratio=2). (B) Mean performance of HESTIA across seven samples under different weights of the cross-resolution alignment loss. Curves show the mean values of ARI, NMI, ASW, PAS, and CHAOS, and shaded areas indicate one standard deviation across samples. The vertical dashed line marks the benchmark setting (α=1.0).

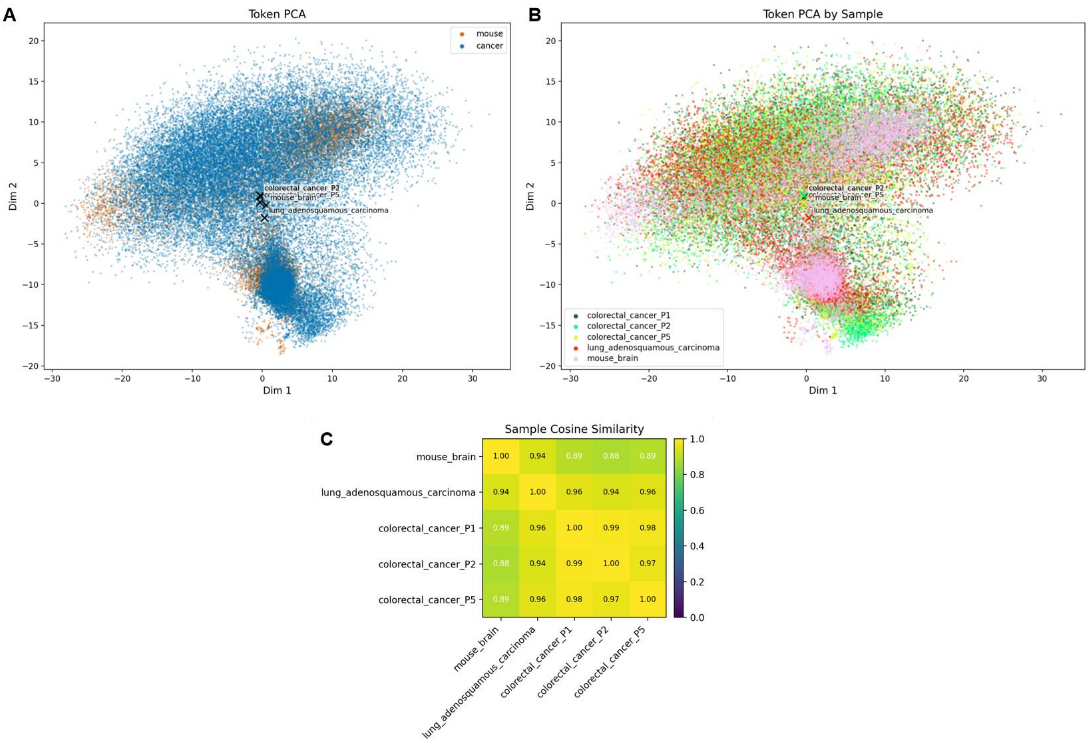


**Supplementary Figure 12 Assessment of HIPT embedding consistency between mouse brain and cancer H&E samples.**

(A) Token-level PCA of 10,000 randomly sampled spatial tokens per sample. The silhouette score is about -0.002. (B) The same token-level PCA colored by sample identity. (C) Heatmap of cosine similarity between sample-level pooled HIPT embeddings.

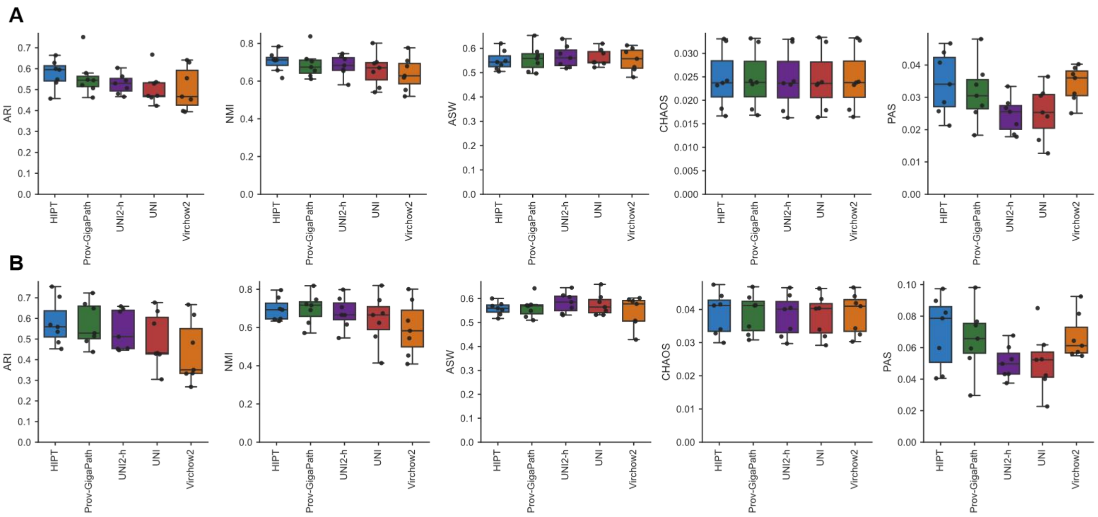


**Supplementary Figure 13 Performance of HESTIA with alternative histology feature extractors.** Boxplots showing ARI, NMI, ASW, CHAOS, and PAS for HESTIA with HIPT, Prov-GigaPath, UNI, UNI2-h and Virchow2 across seven mouse brain subregions at bin20 level (A), and cellbin level (B). All HESTIA variants were trained with default hyperparameters without specific tuning.
